# De-risking drug discovery of intracellular targeting peptides: screening strategies to eliminate false-positive hits

**DOI:** 10.1101/636563

**Authors:** Simon Ng, Yu-Chi Juang, Arun Chandramohan, Hung Yi Kristal Kaan, Ahmad Sadruddin, Tsz Ying Yuen, Fernando J. Ferrer, Xue’Er Cheryl Lee, Liew Xi, Charles W. Johannes, Christopher J. Brown, Srinivasaraghavan Kannan, Pietro G. Aronica, Nils Berglund, Chandra S. Verma, Lijuan Liu, Alexander Stoeck, Tomi K. Sawyer, Anthony W. Partridge, David P. Lane

**Author notes:** **Footnote** Author contributions: S.N., Y.J., A.C., H.Y.K.K., A.S., T.Y.Y., F.J.F., C.W.J., C.J.B., L.L., A.S., A.W.P. designed research; S.N., Y.J., A.C., H.Y.K.K., A.S., T.Y.Y., F.J.F., X.C.L., L.X., L.L., A.S., S.K., P.G.A., N.B. performed research; S.N., Y.J., A.C., C.W.J., C.J.B., C.S.V., A.S., A.W.P. analyzed data; S.N., A.C., C.J.B., C.W.J., S.K., T.K.S., A.W.P., D.P.L. wrote the paper.

## Abstract

Discovery of false-positive target binding, due to assay interference or aggregation, presents a significant problem for drug discovery programs. These issues may often be unrealized and could lead researchers astray if not subject to independent verification of reproducibility and/or on-target mechanism of action. Although well-documented for small molecules, this issue has not been widely explored for peptide modality. As a case study, we demonstrate that two purported KRas inhibitors, stapled peptide SAH-SOS1*_A_* and macrocyclic peptide cyclorasin 9A5, exemplify false-positive molecules – both in terms of their sub-micromolar KRas binding affinities and their on-target cellular activities. We observed that the apparent binding of fluorescein-labeled SAH-SOS1*_A_* given by a fluorescence polarization assay is sensitive to detergent. False-positive readouts can arise from peptide adsorption to the surface of microplates. Hence, we used surface plasmon resonance and isothermal titration calorimetry to unambiguously show that both SAH-SOS1*_A_* and cyclorasin 9A5 are non-binders for KRas. Thermal shift assay and hydrogen-deuterium exchange mass spectrometry further demonstrate that both peptides destabilize KRas and induce unfolding of the protein. Furthermore, both peptides caused significant release of intracellular lactate dehydrogenase, suggesting that membrane rupture rather than on-target activity is accountable for their reported cytotoxicity. Finally, both peptides exhibited off-target activities by inhibiting the proliferation of U-2 OS and A549 cells, despite their independency of the KRas signaling pathway. Our findings demonstrate the critical need to employ orthogonal binding assays and cellular counter-screens to de-risk false-positive molecules. More rigorous workflows should lead to improved data and help obviate inadvertent scientific conclusions.

**Significance statement:** False positive molecule hits occur frequently in high-throughput screens and can contaminate the scientific literature. This has become an increasingly serious issue in small molecule drug discovery and chemical probe development and it is not surprising that peptides may be similarly prone to assay interference. Using KRas as a target and two known macrocyclic peptide inhibitors as a case study, we clearly show that reporter-free biophysical assays and cellular counter-screens offer the solution to detect and de-risk the potential of false-positive compounds. We further discuss the advantages, limitations and overall strategic importance of such methods.

## Introduction

Drug discovery programs are often initiated with efforts aimed at identifying molecules that bind their targets with high affinity to modulate biological activity with high selectivity and potency. Ensuring that the binding interaction is authentic is crucial as the risk of being misled by a false positive is substantial. Indeed, examples show that misleading biological activities given by questionable compounds (1, 2), if left unchecked, can propagate in the scientific literature (3, 4). Consequently, there may be undesirable effects within both research and drug discovery in terms of time and investment to pursue molecules that are not true benchmarks to further investigate target-driven disease mechanisms (5–7)

Pan-assay interference (8) and colloidal aggregation (9) may underscore the basis for most frequently identified false-positive molecules. Increasing evidences have emerged in the literature to elucidate such promiscuous activities (5, 10, 11). These non-specific and misleading activities could occur through several mechanisms (10) which include covalent reactivity, redox cycling, fluorescence interference, membrane disruption, and the formation of colloidal aggregates. Among them, colloidal aggregate formation is perhaps the most common (12, 13). In seminal work (9), Shoichet and colleagues proposed that the colloidal aggregates exert their effects by enzyme sequestration, thereby blocking and inhibiting their activities in unexpected ways. The lists of aggregators are not just limited to synthetic small molecules, and are also found among natural products (14) and marketed drugs (15); these issues are widespread. Besides enzymes, targets involved in protein-protein interactions are affected as well (16, 17). Importantly, these artefacts are difficult to discern if the readout from the assay is reproducible and dose-dependent. One tool for addressing this requires the addition of detergents into the biochemical assay such that the apparent activities of colloidal aggregates can be alleviated or even completely abrogated (18).

Macrocyclic peptides represent an exciting chemical modality with the potential to therapeutically address intracellular protein-protein interactions – these represent targets that are most often intractable with a small-molecule modality (19, 20). Indeed, the macrocyclic peptide approach can produce molecules that exhibit highly specific one-to-one stoichiometric binding and on-target cellular activities (21, 22). In fact, these efforts have translated to a macrocyclic peptide (ALRN-6924) that has entered clinical trials (23). Despite these encouraging developments, we describe herein macrocyclic peptides to also have the potential of effecting false-positive modulation of protein-protein interactions. As a case study, we report evidences that a stapled peptide (SAH-SOS1*_A_*) and a macrocyclic peptide (cyclorasin 9A5), independently described as molecules that bind to KRas and inhibit its downstream pathway, are in fact false-positive molecules, both in terms of their sub-micromolar affinities and their on-target cellular activities (24, 25). We first reproduce the binding isotherm of SAH-SOS1*_A_* using the fluorescence polarization assay reported in the original publication (24). However, to our surprise, adding detergent into the assay solution completely abolished binding. Further experiments suggest that the apparent binding may arise from adherence of the stapled peptide to the microplate. Additionally, using reporter-free techniques, such as surface plasmon resonance and isothermal titration calorimetry, we unambiguously showed that SAH-SOS1*_A_* and cyclorasin 9A5 did not bind to KRas. In thermal shift assay, the melting temperature of KRas decreased in the presence of the peptides as opposed to the no-ligand control, suggesting ligand-induced destabilization of the protein. This phenomenon was further confirmed by the data from hydrogen-deuterium exchange monitored using mass spectrometry. Both peptides induced more deuterium uptake relative to the no-ligand control indicative of destabilization of the protein. Critically, we have validated these biophysical methods for KRas by employing a macrocyclic peptide (KRpep-2d) discovered by Takeda (26) as the positive control in all the assays we used. The same group also delineated the binding of KRpep-2d with a careful study of its structural-activity-relationships (27) and confirmed the binding with a co-crystal structure of KRpep-2d/KRas (28).

In this report, we further showed that the cellular activities of SAH-SOS1*_A_* and cyclorasin 9A5 were attributed to their ability to trigger the lysis of cell membranes at an EC_50_ in the range of 10−30 μM as measured by the release of intracellular lactate dehydrogenase (LDH). The LDH release potencies coincidentally overlap with their reported cytotoxic potencies. Furthermore, both peptides exhibited strong anti-proliferative effects in KRas-independent cell lines, suggesting KRas is not their biological target. Collectively, we conclude that the apparent biological activities reported in both publications may arise from off-target activities and/or non-specific cell death due to the rupture of cell membranes.

## Results and Discussion

### FAM-SAH-SOS1*_A_* is a Promiscuous Binder

Mutant KRas is a significant oncogenic driver (29, 30). Yet, KRas is typically considered a challenging therapeutic target as the surfaces that mediate the interactions between KRas and its downstream effectors lack deep hydrophobic pockets to which a small molecule might bind (31, 32). As an alternative approach, data has supported macrocyclic peptides as a promising modality for targeting protein-protein interactions (19, 20, 33). This prompted us to investigate two widely cited publications published in 2015 by Walensky and colleagues (24) and Pei and colleagues (25). To disrupt the KRas−SOS1 interaction, Walensky and colleagues designed an 18-residue stapled peptide, named SAH-SOS1*_A_*, by incorporating two extra arginines at the N-terminus and an *i*, *i*+4 hydrocarbon linker at the non-interacting face of the SOS1 α-helix (929–944). Using fluorescence polarization (FP) as the primary assay, the investigators showed that a fluorescein-labeled SAH-SOS1*_A_* could bind to wild-type, as well as G12D, G12C, G12V, G12S and Q61H mutated forms of KRas with apparent dissociation constants in the range of 100 to 175 nM. Their results further suggested that SAH-SOS1*_A_* engaged both GDP- and GTP-loaded wild-type KRas with indistinguishable affinities, despite the fact that these two forms of the protein are well known to adopt vastly different conformations in solution (34).

First, we were able to reproduce the binding isotherm of fluorescein-labeled SAH-SOS1*_A_* (FAM-SAH-SOS1*_A_*) using a few selected forms of KRas (G12D, Q61H, and wild-type) loaded with GDP (Fig. 1*A*, Fig. S1). The FP binding curve adopted a sigmoidal shape and seemed saturated, despite the sub-optimal fitting (R^2^ = 0.812, Fig. 1*A*). To our surprise, the apparent bindings were completely abolished upon addition of non-ionic detergent (0.01% v/v Tween 20) to the assay buffer. To check for promiscuous binding, we titrated sequence-unrelated proteins, e.g., MDM2 (Fig. 1*B*) and eIF4E (Fig. 1*C*) separately, into FAM-SAH-SOS1*_A_*. Further confirming the non-specific nature of the signal, FAM-SAH-SOS1*_A_* demonstrated apparent binding to each of these unrelated target proteins with affinities similar to that obtained for KRas. Similarly, these binding interactions were found to be detergent-sensitive. We also tested the negative control peptide (FAM-SAH-SOS1*_B_*), which has a hydrocarbon linker installed at the interacting face of the helix. FAM-SAH-SOS1*_B_* showed no binding in the FP assay as described in the publication (24). However, we found that FAM-SAH-SOS1*_B_* could bind to KRas as well as the unrelated proteins, albeit the signal increases were lower in magnitude (Fig. S2). Again, the non-specific interaction could be abrogated by adding detergent. On the other hand, fluorescein-labeled ATSP-7041 (A^8^Q, Q^9^L), a progenitor of the first stapled peptide entering clinical trials (23), could bind to its target MDM2 and retain the binding in the presence of detergent, suggesting ATSP-7041 (A^8^Q, Q^9^L) is a *bona fide* binder for MDM2 (Fig. S2). In contrast, FAM-SAH-SOS1*_A_* and FAM-SAH-SOS1*_B_* bind to all proteins we have investigated, and the binding interactions are always partially or completely abolished by detergent.

**Fig. 1.**
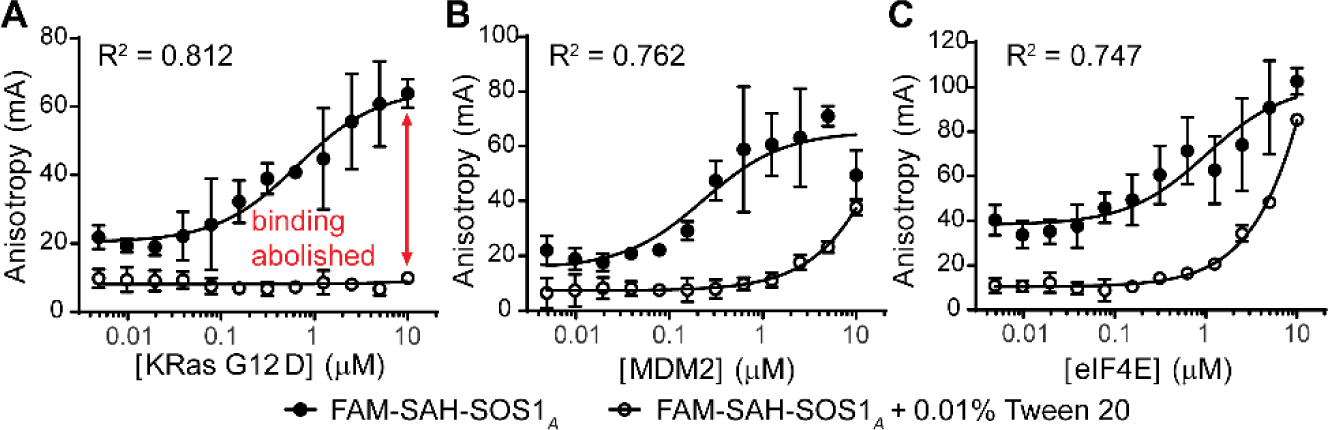
Fluorescence polarization assays were performed in uncoated black polypropylene microplate using FAM-SAH-SOS1*_A_* (15 nM) and varying concentration of three distinct protein targets. The peptide demonstrated binding to (*A*) KRas G12D, (B) MDM2, and (C) eIF4E. However, the bindings were abolished or attenuated in the presence of detergent. Data are mean of technical triplicates ± SD. R^2^ corresponds to the fitting of no-detergent data (solid circle).

### Mechanisms for the False-positive Readout in Fluorescence Polarization

We hypothesize that, in the absence of detergent, two scenarios could contribute to the false-positive FP signals: (i) FAM-labeled peptide non-specifically adsorbs on to the plastic surface of the microplate, and/or (ii) FAM-labeled peptide forms a colloidal aggregate which then sequesters the proteins. Under both circumstances, the apparent “hydrodynamic size” of the peptide increases, leading to a slower tumbling rate and an increased fraction of polarized light. As a result, the FP signals increase and give a false impression that the peptide binds, one-to-one, to a protein.

To test whether the microplate could influence the assay, we explored three different types of microplates: (i) uncoated polypropylene microplate (used for the experiments in Fig. 1 and Fig. S1 and S2); (ii) uncoated polystyrene microplate; and (iii) polystyrene microplate coated with a non-ionic hydrophilic material (Corning NBS^TM^). We observed apparent binding of FAM-SAH-SOS1*_A_* regardless of the use of polystyrene or polypropylene microplate (Fig. 2*A* and 2*B*). The apparent binding disappears once we added the detergent or switched to a polystyrene microplate with a non-binding surface (Fig. 2*C*). Thus, it appeared that the increased FP signal was related to non-specific adherence of FAM-SAH-SOS1*_A_* to the plastic surface or protein-coated plastic surface rather than a specific biomolecular interaction with KRas. Indeed, the degree of absorption is minimal for a microplate of hydrophilic nature, with or without the detergent (Fig. 2*C*). Although the amount of KRas protein added to the sample did increase the magnitude of the FP signal, we are confident that this effect is non-specific. This is backed by two observations. First, addition of non-ionic detergent abrogated the FP signal, presumably by preventing non-specific interactions between the peptide and the protein, plastic surface, or protein-coated plastic surface. Second, no binding was observed in the coated microplate (Corning NBS^TM^), regardless of protein concentration. Furthermore, we found that the adsorption of FAM-SAH-SOS1*_A_* to the uncoated microplate is time-dependent. This is supported by a global increase of the FP signals when we delayed the signal measurement by including a pre-incubation time (Fig. S3). To rule out the possibility that KRas was contributing to the signal increase, we monitored the FP signals of FAM-SAH-SOS1*_A_* alone over a course of three hours (Fig. S4). The signal should remain flat in the absence of protein. However, we observed time-dependent signal increase, a phenomenon that likely occurs due to the adsorption of peptide on to plastic surfaces. Furthermore, the adsorption kinetics is greater in an uncoated polystyrene microplate (Fig. S4*A*) than in an uncoated polypropylene microplate (Fig. S4*B*). This is not surprising because the polystyrene surface is generally more hydrophobic than the polypropylene surface (35). In contrast, the FP signal remained flat for the coated polystyrene microplate (Corning NBS^TM^) regardless of the incubation time or the concentration of peptide used (Fig. S4*C*). All data above, thus far, support our hypothesis that adsorption of FAM-labeled peptide to the plastic surface is responsible for the apparent binding observed in the FP assays.

**Fig. 2.**
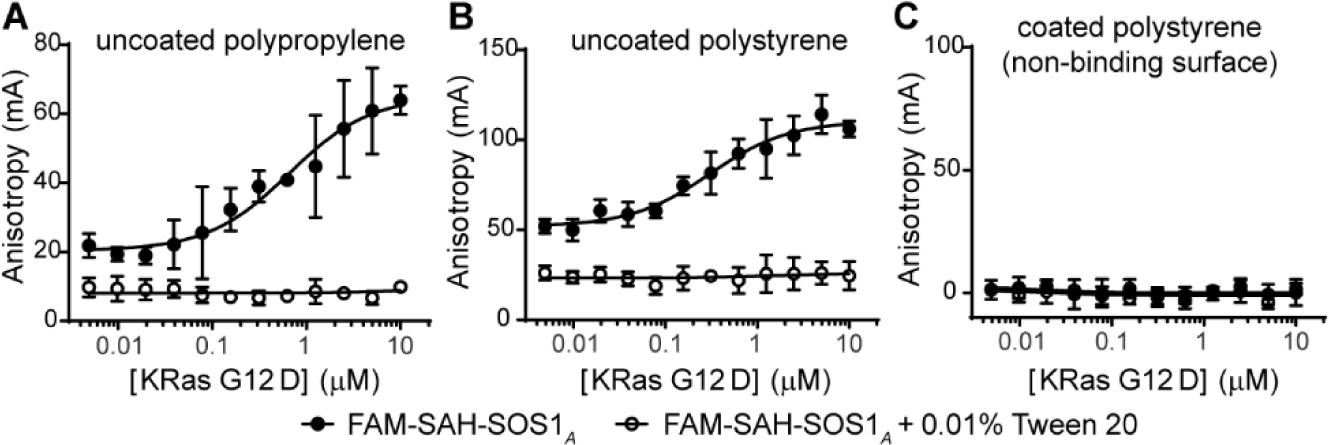
The material of microplate affected the fluorescence polarization of FAM-SAH-SOS1*_A_* (15 nM) and KRas G12D. (*A*) Uncoated polypropylene and (*B*) uncoated polystyrene microplate are hydrophobic and probably adsorbed the peptide and protein in the absence of detergent. (*C*) In contrast, the apparent binding of the FAM-SAH-SOS1*_A_* to KRas disappeared, when we used a polystyrene microplate coated with a hydrophilic material (Corning NBS^TM^). Data are mean of technical triplicates ± SD.

### Detection of Aggregators by Dynamic Light Scattering

Alternatively, FAM-labeled peptides may form colloidal aggregates which can sequester proteins, therefore giving a false-positive FP readout. Assay interference due to aggregation is usually time- and concentration-dependent (9), and the readouts are sensitive to detergent (18). To characterize the peptides, we employed dynamic light scattering (DLS) as a universal technique to directly monitor aggregation. We observed strong and well-defined autocorrelation functions which suggest aggregation in 10 μM solution of FAM-SAH-SOS1*_A_* and FAM-SAH-SOS1*_B_* (Fig. 3). Both peptides gave strong scattering intensities (5700−7400 kcps for FAM-SAH-SOS1*_A_* and 12600−14500 kcps for FAM-SAH-SOS1*_B_*), whereas the signal is much lower for the buffer alone (35−36 kcps). The hydrodynamic diameters of the particles range from 350 to 370 nm for FAM-SAH-SOS1*_A_* and 1200 to 1600 nm for FAM-SAH-SOS1*_B_*. However, we could not detect reproducible and well-defined DLS signals for SAH-SOS1*_A_*, SAH-SOS1*_B_* and cyclorasin 9A5 (data not shown). There are either no aggregates or the aggregates are not stable for these label-free peptides at the concentration tested (10 μM). Interestingly, the ATSP-7041 (A^8^Q, Q^9^L) showed aggregation at 10 μM (Fig. S5), yet it was a specific inhibitor of MDM2 when used at a much lower concentration (<100 nM, Fig. 4). It is noted that, at the tested concentration of 10 μM, it is insufficient to conclude that aggregates are responsible for the false-positive FP readout. We conclude that this most likely derives from direct binding of the peptide to the plastic surface of the microplate.

**Fig. 3.**
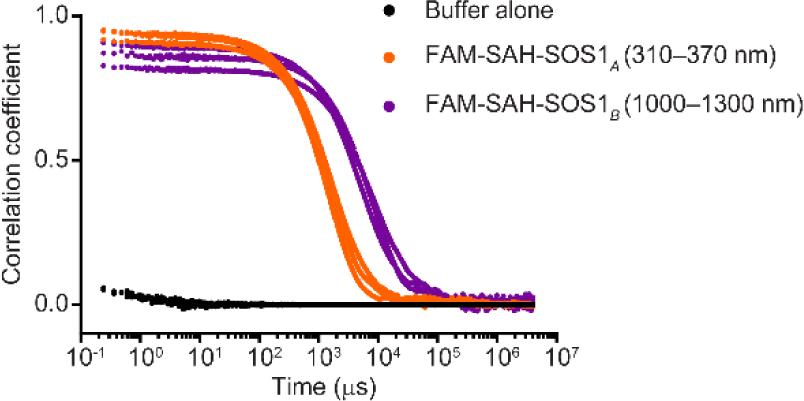
Normalized autocorrelation functions of dynamic light scattering. Each peptide (10 μM) underwent three independent measurements at 25 °C. All data are plotted (range of the particle size in bracket).

**Fig. 4.**
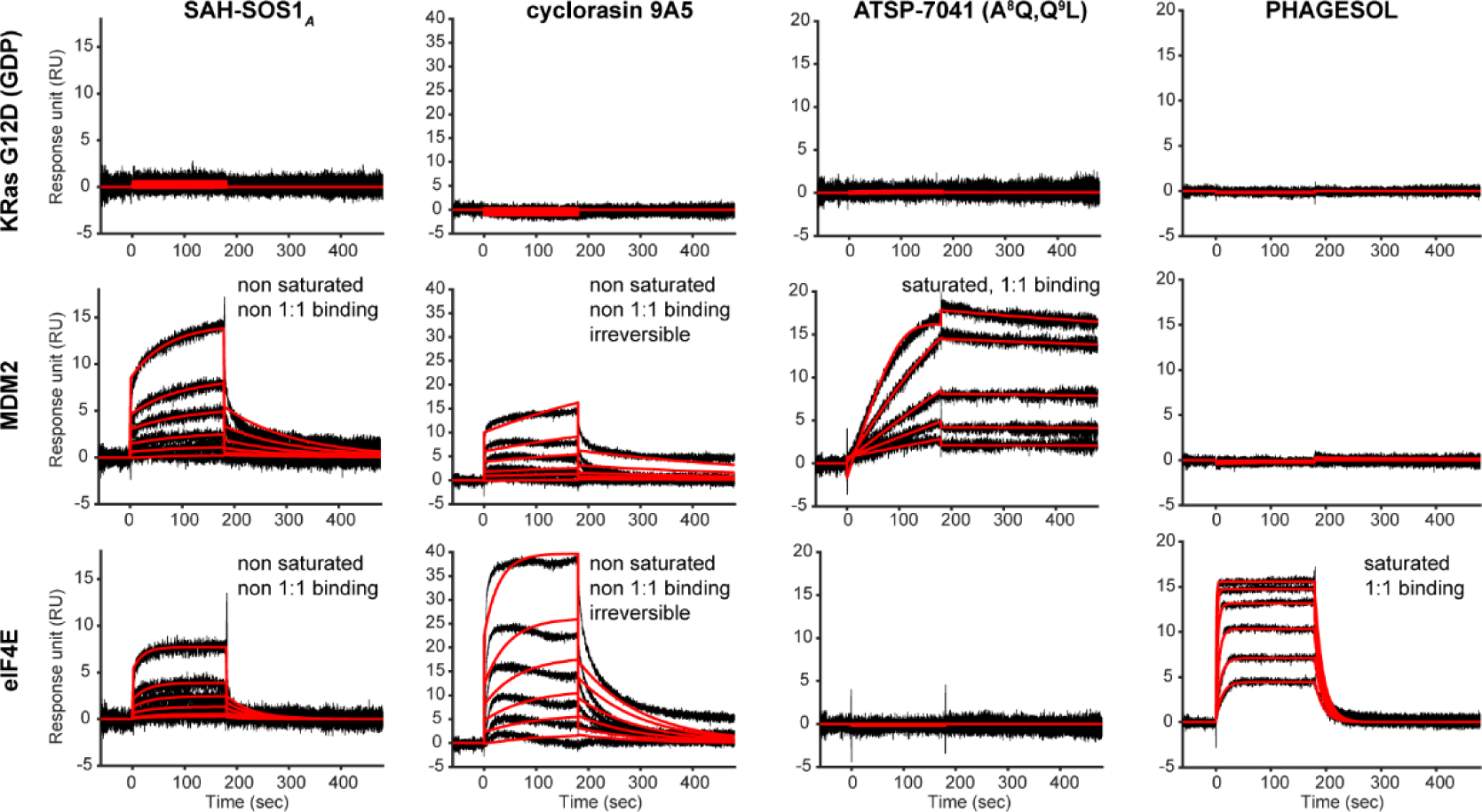
Surface plasmon resonance. We injected each peptide (in column) over the individual flow cell (in row) immobilized with KRas (∼400 RU), MDM2 (∼1200 RU) and eIF4E (∼1600 RU). SAH-SOS1*_A_* (1 μM from top, 2× dilution) and cyclorasin 9A5 (1 μM from top, 2× dilution), did not bind to KRas but demonstrated promiscuous binding to MDM2 and eIF4E. Whereas, ATSP-7041 (A^8^Q, Q^9^L) (50 nM from top, 2× dilution) and PHAGESOL (300 nM from top, 2× dilution) demonstrated saturated 1:1 binding for their corresponding protein targets. Black lines depict the double-referenced sensograms; red lines depict the global fit of the data to a 1:1 binding model.

### Surface Plasmon Resonance Confirms SAH-SOS1*_A_* and Cyclorasin 9A5 as Non-binders for KRas

To carry out a more rigorous analysis, we perceived that it was crucial to run the putative peptides through an orthogonal binding assay, ideally an alternative reporter or reporter-free technique, to ensure that the activity measured by the primary assay is genuine and target-dependent. As a result, we employed surface plasmon resonance (SPR) as the first-pass orthogonal assay. SPR is a reporter-free technique that monitors the association and dissociation of biomolecules in real time. It provides insights into the stoichiometry and reversibility of the binding across a range of concentrations. Consistent with our results from the FP assays, we found no binding at all for SAH-SOS1*_A_* and cyclorasin 9A5, up to 1 μM, across the flow cell containing KRas G12D (Fig. 4, top row). These results implied that either their dissociation constants (K_D_) are much greater than 1 μM or they simply are non-binders. Similarly, FAM-SAH-SOS1*_A_* also did not bind to KRas G12D as demonstrated by SPR (Fig. S6). To ensure that KRas immobilized on the biosensor is competently-folded and functional, we used KRpep-2d, a macrocyclic peptide discovered by Takeda through phage display (26), as the positive control. Indeed, KRpep-2d could bind to KRas G12D and the binding responses are saturable (Fig. S7). Paradoxically, we observed significant SPR readouts from the flow cells immobilized with MDM2 or eIF4E, when we tested SAH-SOS1*_A_* (Fig. 4), cyclorasin 9A5 (Fig. 4) or FAM-SAH-SOS1*_A_* (Fig. S6). We note that their SPR binding responses did not fit well to the 1:1 binding model. Furthermore, at the highest concentration (1 μM) tested, the apparent binding of cyclorasin 9A5 is functionally irreversible, i.e., it exhibits extremely slow dissociation from the surface of the biosensor.

It is unclear how SAH-SOS1*_A_* and cyclorasin 9A5 interact with MDM2 and eIF4E. Previously, SPR has been demonstrated to recognize aggregates quite early in the discovery process (36). It is rather sensitive to non-specific interactions. Adsorption of the aggregates on to the surfaces of biosensors, even though small in amount, can result in non-stoichiometric binding and amplify the SPR signal in an unexpected way (36). We postulate that the non-saturable SPR binding responses plus the high bulk shifts probably result from the non-specific adsorption of SAH-SOS1*_A_* and cyclorasin 9A5 to the surface-loaded proteins. In contrast, ATSP-7041 (A^8^Q, Q^9^L) binds only to MDM2 but not KRas and eIF4E (Fig. 4). PHAGESOL, a peptide we discovered previously through phage display (37), binds exclusively to its target eIF4E but not KRas and MDM2 (Fig. 4). Critically, both ATSP-7041 (A^8^Q, Q^9^L) and PHAGESOL displayed 1:1 saturated binding.

### Isothermal Titration Calorimetry and Thermal Shift Assay Further Confirms SAH-SOS1*_A_* and Cyclorasin 9A5 as Non-binders for KRas

To cross-validate our data in FP and SPR, we included isothermal titration calorimetry (ITC) as the final orthogonal assay. ITC is one of the most rigorous techniques available for the direct measurement of biomolecular interactions under homogenous conditions. Unlike FP or SPR, ITC does not require a fluorescent label or immobilization of the protein/ligand, therefore making it a truly label-free technique. Critically, ITC also directly measures the stoichiometry of binding which makes it ideal to distinguish true binders from artefacts. The drawback, however, is that ITC is relatively low-throughput and demands high amounts of protein (μg-mg) per peptide per run. Nevertheless, we confirmed that SAH-SOS1*_A_* and cyclorasin 9A5 did not bind to KRas G12D using ITC (Fig. 5*A* and Fig. 5*C*). The heat released during the titration of SAH-SOS1*_A_* to the protein or to the buffer itself are almost identical (Fig. 5*A* and Fig. 5*B*). Importantly, cyclorasin 9A5 did not bind to KRas G12V which was the protein reported for the original discovery of the macrocyclic peptide (Fig. 5D). To demonstrate that the KRas used for ITC is well-folded, we showed that the positive control (KRpep-2d) binds to KRas G12D with a K_D_ of 21 nM and a stoichiometry close to unity (Fig. 5*E,* stoichiometry of 0.625 indicates not all the KRas protein is well folded). It is well known that wild-type KRas adopts distinct conformations and functions when bound to different nucleotides. As a confirmatory experiment, we show that wild-type KRas binds to its downstream effector (Raf-RBD) only when KRas is activated with GMPPNP (K_D_ = 6 nM, N = 0.554) but not with GDP (Fig. 5*F* and 5*G*).

**Fig. 5.**
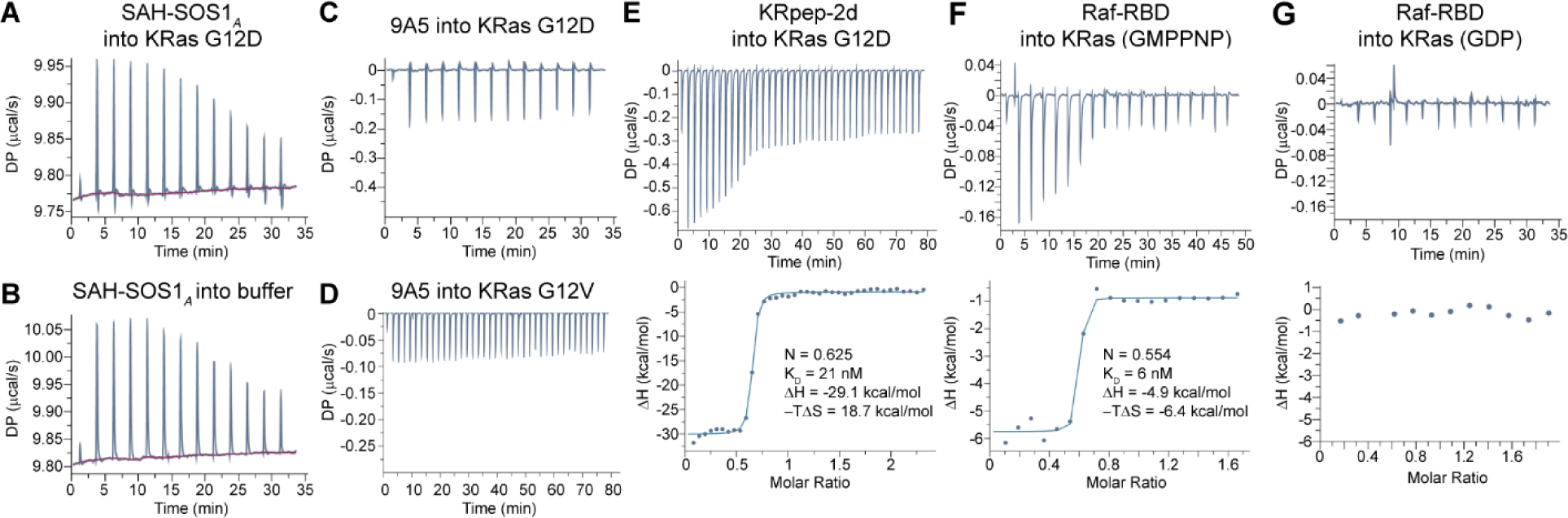
Isothermal titration calorimetry. (*A*) The heat changes from titrating SAH-SOS1*_A_* into KRas G12D. (*B*) The heat changes from titrating SAH-SOS1*_A_* into buffer alone. (*C*) The heat changes from titrating cyclorasin 9A5 into KRas G12D or (*D*) KRas G12V. (*E*) KRpep-2d demonstrated tight binding to KRas G12D. (*F*) Raf-RBD demonstrated tight binding to activated wild-type KRas (GMPPNP) but no binding to the (*G*) inactive wild-type KRas loaded with GDP.

We also performed thermal shift assays for SAH-SOS1*_A_* and cyclorasin 9A5. Both peptides produced a negative shift in the melting temperatures (ΔT_m_) of KRas G12D (Fig. S8), as compared to the no-ligand control, indicating ligand-induced destabilization. At higher concentrations, SAH-SOS1*_A_* displayed high initial background probably due to aggregated peptide-dye interactions (Fig. S8*A*). Furthermore, a global service partner (Proteros) also failed to detect binding of either peptide with crystal-grade protein using ITC and thermal shift assay (data not shown). As an important positive control, KRpep-2d was able to stabilize KRas with a significant positive ΔT_m_ of +11.5 °C (Fig. S8*C*), indicating a well-folded protein capable of binding to specific ligands. To ensure our biophysical observations were not unique to cyclorasin 9A5, we tested a closely related analog, cyclorasin 9A16, a molecule with reported IC_50_ values that were similar to cyclorasin 9A5’s in both the inhibition of KRas-Raf interaction and anti-proliferative assay (25). Once again, the thermal shift assay produced a negative ΔT_m_ shift (−5.9 °C) for cyclorasin 9A16 (Fig. S9*A*). By employing SPR and ITC experiments, we further validated cyclorasin 9A16 as a non-binder, i.e., false positive, for KRas (Fig. S9*B* and S9*C*).

### HDX-MS Confirms SAH-SOS1*_A_* and Cyclorasins Destablize KRas

A host of biophysical techniques employed here (FP, ITC, SPR, and thermal shift assay) collectively establish SAH-SOS1*_A_* and cyclorasins (9A5 and 9A16) as false positive KRas binders. Additionally, the negative ΔT_m_ shifts in the thermal shift assay suggest these molecules non-specifically destabilize KRas. To confirm these observations and to capture protein-wide effects of the three peptides (SAH-SOS1*_A_*, cyclorasin 9A5 and cyclorasin 9A16) on KRas G12D, hydrogen-deuterium exchange mass spectrometry (HDX-MS) was used to monitor changes in deuterium uptake across the protein amides in response to the binding of the peptides. HDX-MS experiments were carried out to compare deuterium uptake between unbound-KRas G12D (GDP) and peptide-bound KRas G12D. Briefly, KRas G12D in the presence and absence of individual peptides (SAH-SOS1*_A_*, cyclorasin 9A5 and cyclorasin 9A16) was labelled with deuterium for 1 and 5 min each. The labelled protein was proteolyzed into peptide fragments and the deuterium uptake of each fragment was quantified by mass spectrometry in the presence and absence of the peptides. HDX-MS experiments of KRas identified 88 pepsin-proteolyzed peptides with primary sequence coverage of 100 % (Fig. S10). Complete sequence coverage allowed us to assess binding or destabilization across the entire protein sequence.

Application of HDX-MS to various peptide/KRas G12D combinations was initiated with employment of a variant of KRpep-2d, without terminal arginines (KRpep-2d (Arg del), Fig. S7), as a positive control, due to its better solubility compared to the parent KRpep-2d. The corresponding data gave a HDX protection pattern highly consistent with the binding site observed in the KRas/KRpep-2d co-crystal structure (28), thus validating the technique’s ability to faithfully map the peptide-binding sites (Fig 6). In contrast, the experiments involving SAH-SOS1*_A_*, cyclorasin (9A5 and 9A16) demonstrated a complete absence of specific binding and agreed well with the destabilization (Fig. 6E) observed in the thermal shift assay (Fig. S8 and S9A). Specifically, the HDX-MS data suggest non-specific and protein-wide destabilization of KRas G12D in the presence of SAH-SOS1*_A_*, cyclorasin (9A5 and 9A16) (Fig 6A−C). All three peptides showed similar deuterium uptake profiles suggestive of similar effects on KRAS G12D. In addition, no significant binding-induced protection from deuterium uptake was observed at any pepsin-proteolyzed KRas G12D peptides, which confirms the absence of binding of SAH-SOS1*_A_*, cyclorasin (9A5 and 9A16) to KRas G12D. HDX experiments monitoring binding of positive control peptide to KRas G12D showed protection from deuterium uptake at regions previously identified as binding site for KRpep-2d from X-ray structural studies (Fig. 6D and 6F) (28). Together, these results confirm that solution dynamic analysis (using deuterium labeling followed by MS) of SAH-SOS1A, cyclorasin (9A5 and 9A16) show global destabilization of KRAS G12D, unlike positive control KRpep-2d (Arg del), where binding-induced protections of specific protein’s regions were observed.

**Fig. 6.**
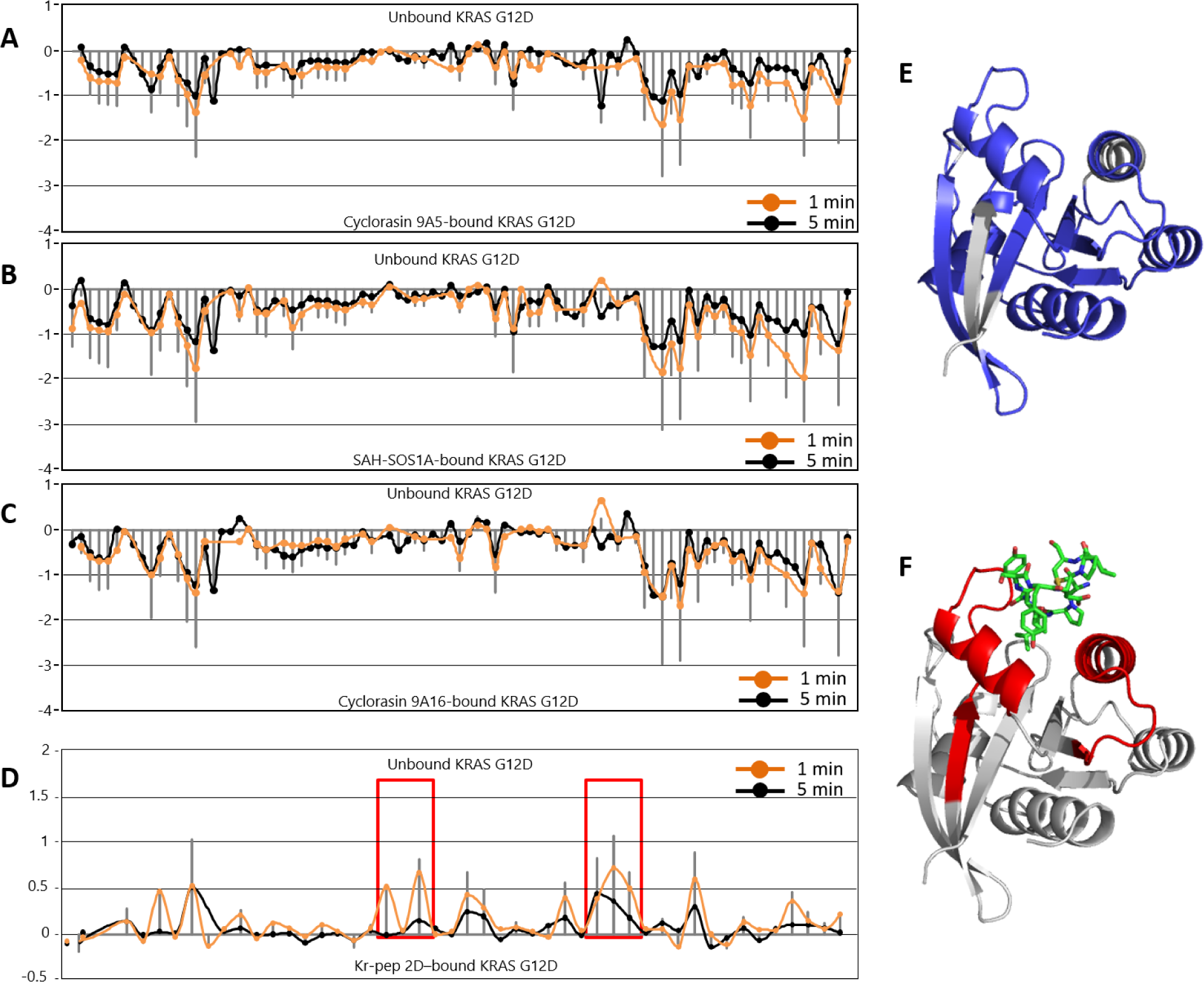
HDX-MS. Deuterium uptake across KRas G12D peptides is plotted as difference plots, comparing the deuterium uptake in the unbound KRas G12D vs. peptide-bound KRas G12D. Differences in deuterium exchange (in Daltons) is plotted on the Y-axis and pepsin-proteolyzed peptides are plotted on the X-axis from N- to C-terminus. Differences in deuterium uptake at each of these peptides are plotted at 1min (orange) and 5 min (black) of deuterium labeling time-points. Each dot represents a single pepsin-proteolyzed peptide from KRas G12D and solid grey lines represent sum of difference at both 1- and 5-min time-points. Positive differences (Y-axis) are suggestive of protection from deuterium uptake in the presence of the peptide and negative differences are indicative of destabilization in the presence of peptide. Difference in deuterium uptake across KRAS G12D peptides in the presence of (*A*) cyclorasin 9A5, (*B*) SAH-SOS1*_A_*, and (*C*) cyclorasin 9A16 show increase in deuterium uptake across multiple regions of KRas G12D suggestive of global destabilization/unfolding. (*D*) Addition of positive control KRpep-2d (Arg del) to KRAS G12D shows decreases in deuterium uptake at the X-ray crystallography-identified binding regions of KRpep-2d (boxed in red). (*E*) Residues with increase in deuterium uptake in the presence of cyclorasin 9A5 are mapped (highlighted in blue) onto the structure of KRas G12D. Residues with no significant differences (<0.5 Da) are colored grey. (*F*) Residues that show protection from deuterium exchange (boxed red in (*D*)) in the presence of KRpep-2d (Arg del) are mapped on to the structure of KRas G12D (PDB: 5XCO). The highlighted regions identified as binding site by HDX (in red) form the 3D binding pocket for KRpep-2d (represented by green stick view) on KRas G12D when mapped onto the structure. Arginines in KRpep-2d are hidden for clarity purpose.

### Modelling and molecular simulations of the putative complexes between KRas and SAH-SOS1*_A_*, Cyclorasin 9A5 or KRpep-2d

A series of peptides inspired by the observations of Walensky and colleagues (24) were designed and docked to the structure of Ras complexed to SOS. These peptides were subject to MD simulations to monitor their stability (Fig. S11). None of the peptides (unstapled and stapled) were found to remain in the initial (and speculated) location and were observed to drift away; this agreed with the experimental observations that none of the peptides bind to KRas. In a similar manner, the complexes between KRas and cyclorasin 9A5 were generated and subject to MD simulations. Once again, the peptides did not bind stably and drifted away (Fig. S12), in agreement with the lack of binding seen in experiments. In contrast, KRpep-2d remained bound to GDP-bound KRas, close to the crystal structure, during the triplicate simulations.

### SAH-SOS1*_A_* and Cyclorasin 9A5 can Disrupt Cell Membranes and Trigger Cell Lysis

With multiple techniques showing that SAH-SOS1*_A_* and cyclorasin 9A5 are false-positive binders of KRas, we were perplexed by the biological activities reported in the publications (24, 25). Release of intracellular lactate dehydrogenase (LDH) is a common method used to directly quantify the degree of cell membrane rupture. We could detect appreciable LDH leakage for SAH-SOS1*_A_* (EC_50_ = 10 μM), SAH-SOS1*_A_* with unstapled hydrocarbon (EC_50_ = 8 μM) and cyclorasin 9A5 (EC_50_ = 30 μM) (Fig. 7). The EC_50_ of LDH leakage for these peptides is comparable to the EC_50_ of the positive control (iDNA79, EC_50_ = 20 μM). iDNA79 is a cationic amphipathic stapled peptide we discovered in-house that gave strong lytic activity. In contrast, the LDH release is insignificant (< 10%) for SAH-SOS1*_B_*, Aib-SOS1*_A_*, cyclorasin 12A, ATSP-7041 (A^8^Q, Q^9^L) and PHAGESOL, and minimal (< 30%) for cyclorasin 9A54, up to 50 μM of the concentration tested.

**Fig. 7.**
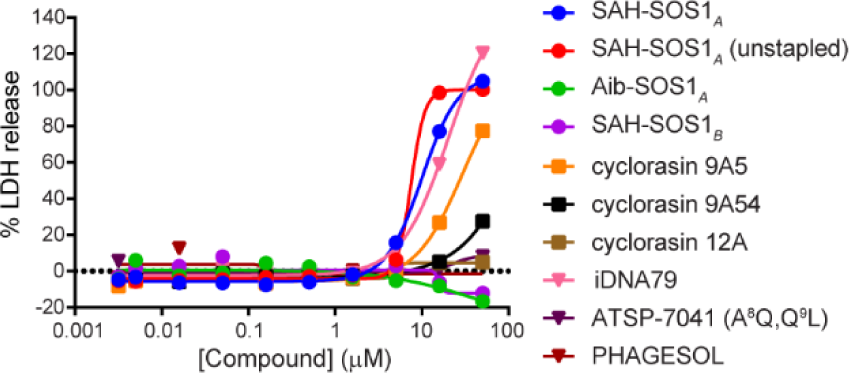
Lactate dehydrogenase (LDH) release assay. We treated HCT116 cells with the peptides under serum-free condition for 4 hours, followed by quantifying the degree of LDH release in the media. Maximum LDH release was defined as the amount of LDH released induced by the lytic peptide (iDNA79) and used to normalize the results.

We performed the LDH release assay using HCT116, a colorectal cell line which harbors mutant KRas^G13D^, under serum-free condition. Although serum could influence the assay, we believe the condition is relevant because the viability assay reported in the publication (24) is performed under serum-free condition during the peptide treatment. We incubated the cells with peptides for 4 hours. We expect that rapid cell death within this short treatment time is unlikely to involve cell cycle arrest and apoptosis due to the inhibition of KRas signaling pathway. The rupture of plasma membrane, a hallmark of non-specific cell death, is most likely the cause. The LDH results suggest that SAH-SOS1*_A_* and cyclorasin 9A5, when used at sufficiently high concentration (10–30 μM), may have compromised the plasma membrane, and caused the lysis and rapid death of cells. Interestingly, these undesired toxicities are only observed for SAH-SOS1*_A_* and its unstapled hydrocarbon analog, but not for its linear version (Aib-SOS1*_A_*), and the negative control (SAH-SOS1*_B_*) where the stapled hydrocarbon is placed at a different face of the helix. Minimal LDH leakage is also observed for the analogs of cyclorasin 9A5, such as cyclorasin 9A54 and cyclorasin 12A. Coincidentally, the degree of membrane disruption caused by the SAH-SOS1*_A_*, cyclorasin 9A5 and their non-toxic analogs is consistent with the cytotoxicity reported in the publications (24, 25). In a different report, Bird *et al.* disclosed that SAH-SOS1*_A_* did not induce LDH release (38). But, we found that the tested peptides are actually different from the original SAH-SOS1*_A_* (24). The investigators changed the length of the hydrocarbon staple and removed the two arginine residues from SAH-SOS1*_A_* that could be accountable for the membrane disruption.

### SAH-SOS1*_A_* and Cyclorasin 9A5 Exhibit Off-target Cytotoxic Activities in KRas-independent Cell Lines

To further investigate the promiscuous cellular activities, we assessed the effects of SAH-SOS1*_A_* and cyclorasin 9A5 on the proliferation of KRas-independent cell lines (U-2 OS cells and A549 cells). These cell lines do not require KRas to maintain viability and their downstream signaling is not responsive to KRas knockdown by gene silencing (39, 40). Strikingly, above a concentration threshold of 20 μM, we observed strong anti-proliferative effects of SAH-SOS1*_A_* and cyclorasin 9A5 on both cell lines (Fig. 8*A*). The results suggest that both peptides act through off-target mechanisms, since inhibition of the KRas signaling pathways should not impede the proliferation of these cells. We also note that the dose-response curves are very steep, i.e., a small increment of peptide concentration (< 6-fold change) is enough to induce a strong anti-proliferative effect (a decrease from plateau to baseline level).

**Fig. 8.**
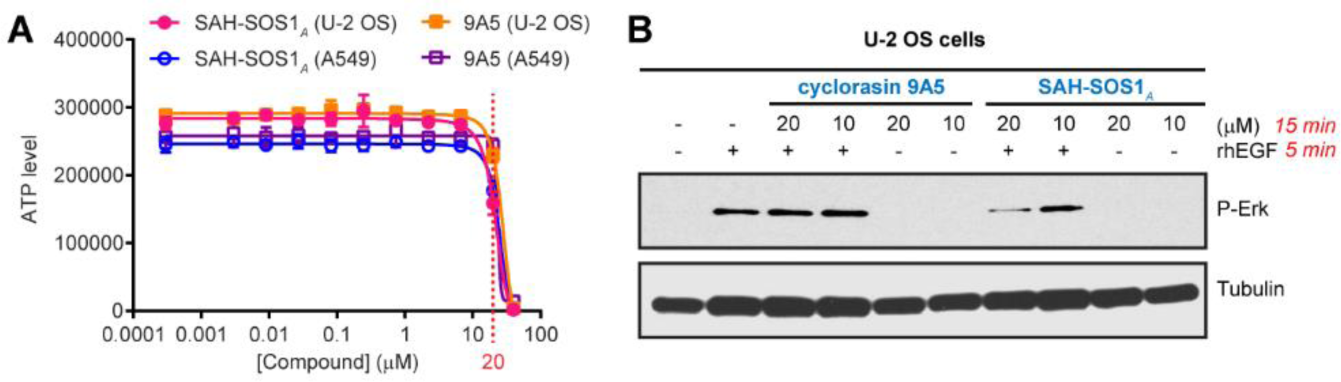
Anti-proliferation and inhibition of the downstream signaling in KRas-mutant (A549) and WT (U-2 OS) cell lines. (*A*) 72 hours treatment with SAH-SOS1*_A_* and cyclorasin 9A5, at a threshold above 20 μM concentration, resulted in strong anti-proliferation in U-2 OS and A549 cells by measuring the intracellular ATP level. Data are mean of technical triplicates ± SD. (*B*) 15 min treatment with SAH-SOS1*_A_* but not cyclorasin 9A5 inhibited the downstream phosphorylation of Erk at 20 μM concentration.

We further investigated the effect of the peptides on the KRas signaling pathways using U-2 OS cells (Fig. 8*B*). We showed that SAH-SOS1*_A_* but not cyclorasin 9A5 could inhibit the phosphorylation of Erk, a KRas downstream target, at 20 μM concentration with just 15 minutes of treatment. This result should be interpreted with caution. It is premature to regard the reduced downstream cellular responses as a confirmation of target engagement. Previously, we showed that SAH-SOS1*_A_* triggered rapid cell lysis at EC_50_ of 10 μM. Therefore, the attenuation of phosphorylated Erk observed here may arise from the lysed cells having disrupted membrane-bound KRas and/or compromised cellular machineries which become unresponsive to stimulation by recombinant human EGF. Consistent with the LDH data, our results for the anti-proliferation of KRas-independent cell lines and reduced KRas signaling support the interpretation of the cell death caused by the peptides as non-specific and off-target.

### Concluding Remarks

FP assay is by far the most popular technique (41) for validating protein-protein interactions. Its advantages of high-throughput, mix-and-read format, and the relatively low demand of proteins, makes its cost lower when compared to other techniques. When using a well-validated probe, competitive FP assay is a powerful primary screen for a large library of compounds. However, we showed that direct binding FP assay could be prone to false-positive results. In this report, we demonstrated that fluorescein-labeled peptides gave a false impression of “binding” in FP assay when detergent is omitted in the assay buffer. This is most likely derived from the adherence of the peptide to the plastic surface or protein-coated plastic surface. Such assay interference could mislead the interpretation and are difficult to discern. By employing SPR and ITC as the reporter-free assays, we unambiguously showed that SAH-SOS1*_A_* and cyclorasin (9A5 and 9A16) do not bind to the mutant KRas; in contrast, the positive control (KRpep-2d) does. These results have been confirmed by both thermal shift assay and HDX-MS, where protein-wide destabilization of KRas G12D was observed in the presence of SAH-SOS1*_A_*, cyclorasin 9A5 and cyclorasin 9A16.

An independent group have reproduced the binding of cyclorasin 9A5 to KRas using microscale thermophoresis (MST) (42). However, we note that the reported raw data are not typical of MST data. Specifically, the fluorescence signal did not reach a steady state after the temperature jump. The unusual signals could be indicative of aggregation and artefactual binding, since MST is an indirect measurement of binding based on the change of the hydration shell of biomolecules.

While investigating the mechanism of the false-positive FP readout, we observed aggregation of FAM-labeled peptides and label-free peptides using DLS and SPR respectively. The mechanism of peptide aggregation is not yet clear, although studies suggest that some hydrocarbon-stapled peptides have a propensity to self-associate as demonstrated by NMR and DLS (43). Lanreotide, an FDA-approved macrocyclic peptide drug, is able to self-assemble into a hydrogel in water (44). Interestingly, this property has been exploited to control the long-acting gradual release of the drug after its subcutaneous injection. Indeed, a compound that aggregates at a higher concentration may have legitimate biological activity when used at a lower concentration. ATSP-7041 (A^8^Q, Q^9^L) is one such example. Aggregation-prone compounds should not be excluded and disqualified as non-binders until proven otherwise.

Finally, we showed that SAH-SOS1*_A_* and cyclorasin 9A5 triggered LDH release by disrupting the cell membrane. Amphipathic peptides with positive charges at one face and hydrophobic patch at the other tend to disrupt cell membranes and cause non-specific cytotoxicity (45, 46). Both SAH-SOS1*_A_* and cyclorasin 9A5 contain excessive hydrophobic moiety and multiple positively charged residues. However, such observations cannot be generalized as the closely related analogs of SAH-SOS1*_A_* and cyclorasin 9A5 did not induce significant LDH release. Moreover, we observed strong anti-proliferative activities of SAH-SOS1*_A_* and cyclorasin 9A5 for KRas-independent cell lines. Collectively, we deduce that SAH-SOS1*_A_* and cyclorasin 9A5 most likely killed cancer cells, through off-target effects by the disruption of the cell membrane, rather than through the intended attenuation of signaling pathways mediated by KRas. More work is required to elucidate these off-target activities, such as their mechanisms and structural-activity-relationships which is beyond the scope of this report (47). Yet, we note that by improving the physicochemical properties of SAH-SOS1*_A_* and cyclorasin 9A5, these molecules may become a template useful for binding to their target and for further cellular studies.

Though this report has focused on KRas, our approaches to characterize the macrocyclic peptides are also applicable to other target classes involved in protein-protein interactions (Fig. 9). Our findings make a strong case for using reporter-free systems such as SPR and ITC, whenever possible, as a secondary assay for binding validation. SPR is very sensitive to non-specific binding. Non-saturable SPR signals could imply aggregation or some arcane activity (36). In such cases, ITC should be employed instead to ensure getting the ideal 1:1 stoichiometric binding. We acknowledge that SPR and ITC are low in throughput. These methods might not be suitable for the primary screen performed on a routine basis. Yet, they should be used more often, at least, for the most promising lead, in order to ensure that binding is genuine and the stoichiometry is as expected. We also suggest LDH release assay as a convenient counter-screen to ensure that the observed biological activities do not arise from membrane disruption. In contrast to cell viability assays or indirect measurement of downstream cellular responses, direct intracellular target engagement assays are better tools to prove that the desired target is actually bound when cells are treated with the compounds (Fig. 9). For instance, CETSA (48) and NanoBRET (49) are increasingly employed to distinguish on-target versus off-target cancer-killing effects.

**Fig. 9.**
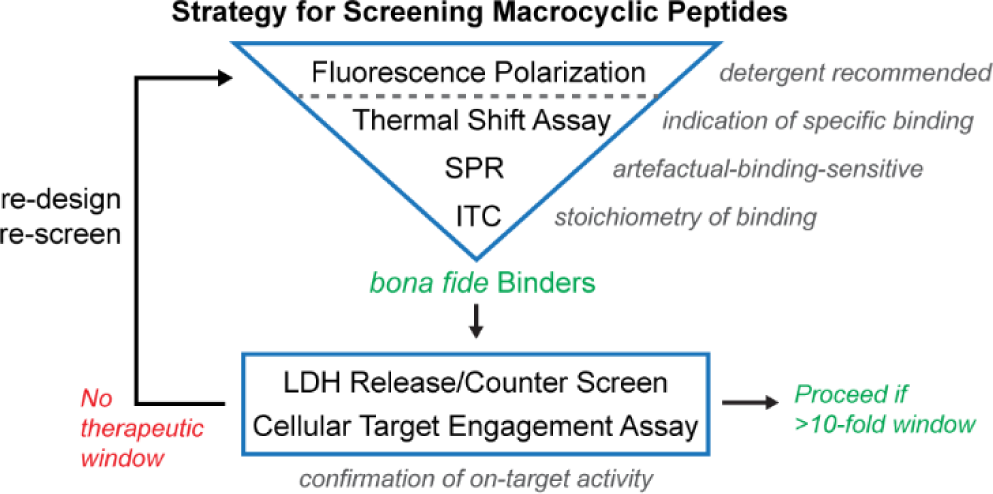
Strategy for screening macrocyclic peptides for intracellular targets to enhance the identification of *bona fide* binders and on-target cellular modulators.

This report documents an extreme case where the molecules investigated do not bind to their target *in vitro*, yet exhibit an apparent cellular activity. In other cases, *bona fide* binders may have confounding cellular activities because of membrane lysis.(50) However, molecules with cellular activity at concentrations well below their membrane lysis threshold remain of interest. Indeed, select examples have been demonstrated to have on-target activity *in vitro* and *in vivo*.(22, 51) *Bona fide* binders with a well-balanced therapeutic window are useful for further studies when applied at a lower dose (Fig. 9). We hope that, by adhering to these principles, it will de-risk assay interference, reveal promiscuous target-unrelated activity, thereby improving the quality of publications to promote a sustainable future for science. Reproducibility does count and orthogonal reporter-free assays should be an essential component of any claim of specific protein-peptide interactions measured by a screening assay based on a fluorescent reporter. Collectively, multiple lines of evidence will serve to strengthen the robustness of the claim.

## Materials and Methods

SAH-SOS1*_A_* and its related analogs were synthesized in house. Cyclorasin 9A5 and its related analogs were sourced from CPC Scientific and IRBM Science Park. ATSP-7041 (A^8^Q, Q^9^L) was synthesized in house. PHAGESOL was synthesized by Mimotopes Pty Ltd. The sequences of the peptides used in this study and their observed m/z could be found in Table S1. All peptides are dissolved in neat DMSO as 10 mM stock solution and diluted thereof for subsequent experiments. Complete descriptions of peptide synthesis, protein expression, biotinylation of protein by sortase ligation, fluorescence polarization assay, dynamic light scattering, surface plasmon resonance, isothermal titration calorimetry, thermal shift assay, hydrogen-deuterium exchange mass spectrometry, molecular simulations, lactate dehydrogenase release assay, cellular growth assay and western blotting are provided in SI Methods.

## Acknowledgements

We thank A*STAR and MSD for the financial support. We also thank Brian Henry for his review of the manuscript draft.

## Supplementary Information Text

### Materials and Methods

#### Peptide synthesis

Peptides were synthesized in-house using the 9-fluorenylmethyloxycarbonyl (Fmoc) solid-phase peptide synthesis (SPPS) method at the 0.1 mmol scale using H-Ramage-Chemmatrix resin (0.53 mmol g^-1^) obtained from PCAS-Biomatrix (Quebec, Canada). All amino acids were purchased from either Advanced Chemtech (Louisville, KY) or Okeanos Tech. Co. Ltd (China). All other solvents and reagents were obtained from Advanced Chemtech (Louisville, KY) or Sigma-Aldrich. All amino acids were N-Fmoc protected and side chains were protected with Boc (Lys, Trp); OtBu (Ser, Thr); Trt (Cys, Asn, Gln). The peptides were synthesized either manually or through the use of a microwave assisted automated peptide synthesizer (CEM-Liberty Blue).

##### Method A: Manual synthesis of peptides

The dry resin was swollen with NMP before use. Deprotection was performed with 20% piperidine in DMF (15 min). Coupling reactions were performed using pre-activated (7 min) solutions of *O*-(7-azabenzotriazol-1-yl)-1,1,3,3-tetramethyluronium hexafluorophosphate (HATU, 3.9 eq), 1-hydroxy-7-azabenzotriazole (HOAt, 4 eq) and DIPEA (8 eq) in NMP (0.5 M). The coupling time was 60 min for all amino acids except for (*S*)-N-Fmoc-2-(4’-pentenyl)alanine and (*R*)-N-Fmoc-2-(7’-octenyl)alanine, which were coupled (3 eq) for 2 hours. After each coupling and deprotection reaction, the resin was thoroughly washed with DMF (5 x 1 min). The amino acids immediately following Fmoc-R8-OH and Fmoc-S5-OH were double coupled.

##### Method B: Automated synthesis of peptides

Deprotection was performed with 20% piperidine in DMF at 90°C for 30 s. Coupling reactions were performed using 5 equivalents each of Fmoc-AA-OH/DIC/Oxyma Pure in DMF (0.2 M) at 90°C for 120 s.

Olefin metathesis reaction was performed on N-terminal Fmoc- or Acetyl-peptides on solid support. The resin was swollen in DCE (pre-dried and degassed) and treated with 6 mM solution of Grubbs’ first-generation catalyst in DCE (2 h). Typically, 3-4 rounds of olefin metathesis treatments were required.

Cleavage of the peptide from the dried resin was achieved using a TFA cocktail consisting of TFA:triisopropylsilane:water (95:2.5:2.5, 8 mL) followed by filtration and precipitation with diethyl ether. The precipitate was collected by centrifugation and dried. The peptide was purified by Reverse Phase High-Performance Liquid Chromatography (RP-HPLC) on GX-271 (Gilson), using a Phenomenex Jupiter C12, 4 μm Proteo 90 Å column (250 x 10 mm). Peptide molecular weight was confirmed by LCMS (Table S1).

#### Protein expression

Full-length human eIF4E was expressed and purified as described previously (1). GGG-MDM2^1-125^ was expressed and purified by NTU protein production platform (Singapore). KRas wild-type, G12D, G12V or Q61H were expressed, purified, and loaded with the desired nucleotide by Evotec AG (Germany).

#### Biotinylation of protein

To attach biotin site-specifically to the protein, we use sortase-mediated ligation. All proteins contain at least one extra glycine at the N-terminus. The ligation was carried out with the protein target (50 μM), sortaseA^61-206/8M^ (1 μM), and biotin-KGGGLPET-GG-OHse(Ac)-amide (200 μM) in 200 μL of ligation buffer (50 mM Tris pH 8.0, 150 mM NaCl, 1 mM TCEP). SortaseA^61-206/8M^ contains mutations that increase ligation efficiency (2) and make it calcium-independent (3). The ligation was incubated at room temperature for 4 hours. The sortase which contains a C-terminal 6×His-tag was removed with Dynabeads His-Tag (cat# 10104D, Thermo Fisher). The biotinylated protein was dialyzed at 4 °C using slide-A-Lyzer cassette (10k MWCO) against 2 L of an appropriate buffer. The buffer was changed after 4−5 hours and the dialysis was repeated for overnight. The biotinylated protein was aliquoted, snap-frozen with liquid nitrogen, and stored at −80°C.

#### Fluorescence polarization assay

All assays were performed at room temperature using assay buffer (50 mM Tris pH 7.4, 150 mM NaCl, 2 mM MgCl_2_ with or without 0.01% v/v Tween 20) and Corning (cat# 3356) 96-well black polypropylene microplate, unless otherwise stated. For experiments performed on uncoated or coated polystyrene microplate, we used Nunc (cat# 237107) 96-well black uncoated polystyrene microplate or Corning (NBS^TM^, cat# 3991) 96-well black coated polystyrene microplate, respectively. The protein was first diluted (2-fold serial dilution) with the assay buffer on the microplate to have a volume of 25 μL in each well. This was followed by the addition of 25 μL of FAM-SAH-SOS1*_A_* (30 nM, dissolved in assay buffer) to each well with multichannel pipette. The final assay solution (50 μL) contains 15 nM of the FAM-SAH-SOS1*_A_* and protein (4.9 nM to 10 μM). Right after the mixing, the microplate was read with EnVision 2104 Multilabel Reader (PerkinElmer) using excitation and emission wavelengths of 480 nm and 535 nm, respectively. The fluorescence anisotropy is plotted against the concentration of protein and the data was fitted with the equation below:

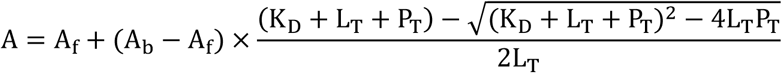

A is the measured anisotropy, A_f_ is the anisotropy for the free peptide (minimum of curve), A_b_ is the anisotropy for the fully bound peptide (maximum of curve), K_D_ is the dissociation constant, L_T_ is the total concentration of the peptide, P_T_ is the total concentration of the protein.

#### Dynamic light scattering

Peptide (10 mM in neat DMSO) was diluted to 10 M with syringe-filtered (0.22 μm) assay buffer (50 mM Tris pH 7.4, 150 mM NaCl, 2 mM MgCl_2_). The solution was analyzed on Zetasizer APS (Malvern Instruments) with 60 mW diode laser (830 nm) at a scattering angle of 90°. The average hydrodynamic diameter of the aggregate represents the intensity-weighted mean derived from a cumulant analysis of the measured correlation curve. Each peptide was analyzed by three independent measurements at 25 °C. Values of the derived count rate (kilo counts per second, kcps) and the average diameter (nm) were reported as a range from the triplicate measurements.

#### Surface plasmon resonance

SPR experiments were performed with Biacore T100 (GE Healthcare) at 25°C. The site-specific mono-biotinylated proteins were prepared by sortase-mediated ligation (see above). SPR buffer consisted of 50 mM Tris pH 7.4, 150 mM NaCl, 2 mM MgCl_2_, 10 μM GDP, 1 mM DTT, 0.01% Tween 20, and 3% DMSO. For SPR study of KRpep-2d and its analog, the reducing agent (DTT) is omitted in the buffer to preserve the disulfide bond in the peptide. The CM5 chip was first conditioned with 100 mM HCl, followed by 0.1% SDS, 50 mM NaOH and then water, all performed twice with 6 sec injection at a flow rate of 100 μL/min. With the flow rate set to 10 μL/min, streptavidin (S4762, Sigma-Aldrich) was immobilized on the conditioned chip through amine coupling as described in the Biacore manual. Excess protein was removed by 30 sec injection of the wash solution (50 mM NaOH + 1 M NaCl) for at least 8 times. The immobilized levels are typically 2500−2800 RU. The biotinylated proteins were captured to the streptavidin, immobilized on independent flow cell, to a level of ∼400 RU for KRas G12D, ∼1200 RU for MDM2, and ∼1600 RU for eIF4E. Flow cell consisted of only streptavidin was used as the reference surface. Using a flow rate of 30 μL/min, peptides dissolved in the SPR buffer are injected for 180 sec. The dissociation was monitored for 300 sec. Each peptide injection is followed by a similar injection of SPR buffer to allow the surface to be fully regenerated. After the run, responses from the target protein surface are transformed by (i) correcting with DMSO calibration curve, (ii) subtracting the responses obtained from the reference surface, and (iii) subtracting the responses of buffer injections from those of peptide injections. The last step is known as double referencing which corrects the systematic artefacts. The resulting responses were subjected to kinetic analysis by global fitting with 1:1 binding model.

#### Isothermal titration calorimetry

The measurements were performed at 25 °C using a Microcal PEAQ-ITC (Malvern Panalytical Inc.). All peptides and proteins were concentrated to 200–300 and 10–20 μM, respectively, in buffer consisted of PBS pH 7.4, 0.03 mM n-dodecyl β-D-maltoside, 1 mM MgCl_2_ and 5% DMSO. Control experiments, which were carried out by injecting peptide into buffer, were subtracted from each data set.

#### Thermal shift assay

The fluorescent dye SYPRO Orange (Invitrogen) was used to monitor thermal denaturation of KRas G12D. Binding of the dye molecule to KRas, as it unfolds due to thermal denaturation, results in an increase in the fluorescence intensity. The midpoint of this transition is termed the T_m_. The thermal shift assay was conducted in a CFX96™ real-time PCR detection system (Bio-Rad). Assay buffer consisted of 50 mM Tris pH 7.4, 150 mM NaCl, 2 mM MgCl_2_ was used for all the dilution. A total 50 μL of mixture containing 5 μL of 31.25× SYPRO Orange (Invitrogen, diluted from 5000× DMSO stock), the peptide of interest, and 10 μM of GDP-loaded KRas G12D was prepared in an 8-well PCR tube. The samples were heated from 25 to 90 °C in 0.5 °C increment each cycle. The holding time for each cycle is 5 sec during which the fluorescence intensity was being measured by Channel 2 (HEX) with Ex/Em:515−535/560−580 nm. Experiments for cyclorasin (100 μM) and KRpep-2d (500 μM) were carried out in duplicate. Experiments for SAH-SOS1*_A_* were carried out in single measurement using varying concentration of the peptide.

#### Hydrogen-deuterium exchange mass spectrometry

Purified KRas G12D protein bound to GDP was incubated with each of the peptides (DMSO for unbound KRas G12D) for a minimum of 30 min to form stable-complexes with KRas G12D concentration of 150 μM, peptide concentration of 500 μM and 5% DMSO in aqueous buffer (10 mM Tris + 1 mM MgCl_2_ pH 7.5). Deuterium exchange experiments were conducted by manually diluting 2 μl of incubated protein-peptide mixture with 48 μl of deuterium buffer (10 mM Tris + 1 mM MgCl2 in D_2_O, pH 7.5) for 1 and 5 min of labelling. Final deuterium labeling concentrations were 6 μM of KRas G12D and 20 μM of peptide-ligands (SAH-SOS1*_A_*, cyclorasin 9A5, cyclorasin 9A16 and KRpep-2d (Arg del)) at 96% D_2_O. 20 μl of 2% TFA-H_2_O quench solution was added for a final pH read of 2.5 to quench the labelling reaction. 50 μl of this quenched reaction was injected into HDX module coupled to an M-class system from Waters Corp. (Milford, MA). All instrumentation, columns, standards and software are from Waters Corp (Milford, MA), unless specified. The sample is passed through Enzymate pepsin column for proteolysis (flow rate of 100 μl/min) and peptides are separated on a C18 acquity analytical column over a 12 min LC gradient (H_2_O vs. ACN at a flowrate of 40 μl/min). The LC-separated peptides are then detected by a Xevo G2-TOF. Data is collected in MSe mode with continuous calibration using glu-fibrinogen peptide (5 μl/min). Peptides are detected using PLGS 3.0 and deuterium uptake is calculated using DynamX 3.0. The deuterium uptake values are averaged from at least two independent HDX-MS experiments.

#### Modelling and molecular simulation

Available crystal structure of KRAS in its apo, GTP (5VQ2) (4) and GDP (4LPK) (5) bound states were used for the generation of the KRAS–peptide complexes. The model of the complex between KRAS and SOS peptide (residues from 929 – 944) complex was generated using the co-crystal structure of HRAS complexed with the SOS protein (1NVU) (6). The staple linkers (all hydrocarbon linkers) were introduced into the SOS peptide to mimic the peptides used by Walensky and colleagues (referred as SAH-SOS1*_A_* in ref (7)), to generate the models of KRAS–SAH-SOS1*_A_* complexes (KRAS was taken in its apo, GTP bound and GDP bound forms). In the absence of a crystal structure of a complex between KRAS and cylcorasin 9A5, we first modelled the 3D structure of the cyclic peptide using the leap module of Amber16 (8) followed by minimization and molecular dynamics (MD) simulations (for 100 ns in triplicates). The conformations of cyclorasin 9A5 generated during the MD simulations were clustered to identify conformational substates of the peptide. Four highly populated conformations were selected for docking to generate models of complexes between KRAS and cyclorasin 9A5. The complexes were generated using blind docking with the program ATTRACT (9). The docked complexes were visually analysed, and the peptides bound at the RBD binding site (as proposed in ref (10)) of KRAS were selected for further MD simulations. The final MD simulations of KRAS (apo, GDP and GTP) complexed to peptides (SOS, SAH-SOS1*_A_* and cyclorasin 9A5) were carried out for 250 ns in triplicates using the program AMBER16 using the standard protocol we have used earlier (11).

#### Lactate dehydrogenase release assay

HCT116 cells were seeded into a 384-well plate at a density of 8000 cells per well. Cells were maintained in McCoy’s 5A Medium with 10% fetal bovine serum (FBS), Blasticidin and Penicillin/Streptomycin. The cells were incubated overnight followed by removal of cell media and addition of Opti-MEM Medium without FBS. Cells were then treated with peptides for 4 hours in Opti-MEM (serum free). Final concentration of DMSO was 0.5%. Lactate dehydrogenase release was detected using CytoTox-ONE Homogenous Membrane Integrity Assay Kit (Promega) as per manufacturer’s instructions. Measurements were carried out using Tecan plate reader. Maximum LDH release was defined as the amount of LDH released induced by the lytic peptide (iDNA79) and used to normalize the results.

#### Cellular growth assay and cell cycle analysis

The A549 and U-2 OS cell lines used in this study were purchased from American Type Culture Collection. Each cell line was maintained in the media and culture conditions recommended by the vendors. There was no evidence of mycoplasma contamination based on a PCR assay. To determine IC_50_ values (growth inhibition as measured by an ATP-based viability assay), cell lines were plated in 96-well plates at 8000 cells per well and treated with increasing concentrations of cyclorasin 9A5 or SAH-SOS1*_A_* and DMSO as vehicle control. Final concentration of DMSO was 0.1%. Cells were plated in triplicates. Viability was determined using the CellTiter-Glo Luminescent Cell Viability assay (Promega, Madison, WI) three days after peptide addition according to the manufacturer’s instructions. Luminescence was measured using a Wallac Victor 3V plate reader.

#### Cell lysis and Western blotting

U-2 OS cells were treated with cyclorasin 9A5 or SAH-SOS1*_A_* for 15 min and stimulated with recombinant EGF at 100 ng/mL for 5 min before it was lysed in lysis buffer (50 mM Tris, pH 8.0, 150 mM NaCl, 1% NP40 supplemented with protease inhibitor cocktail (Thermo Scientific, Rockford, IL)). Protein amounts were determined using the Bio-Rad DC^TM^ Protein Assay Kit II according to the manufacturer’s protocol. Samples were mixed with Laemmli sample buffer (Bio-Rad, Hercules, CA) containing 5% β-mercaptoethanol, separated by 4-15% Tris Glycine extended Gel (Bio-Rad Hercules, CA), and transferred onto a PVDF membrane using Trans-Blot Turbo transfer stacks and the Trans-Blot Turbo transfer apparatus (BioRad, Hercules, CA). The membrane was blocked with 10% non-fat dry milk (BioRad, Hercules, CA) or 5% BSA (Sigma, St. Louis, MO) in TBST (20 mM Tris-HCl, 0.5 M NaCl, 0.1% Tween 20) and incubated with a primary antibody overnight at 4 °C, followed by washes with TBST. Subsequently the membrane was incubated with horseradish peroxidase-conjugated secondary antibody (Jackson Laboratories, West Grove, PA) and detected with ECL developing solution (Thermo Scientific, Rockford, IL). Western blot was performed using phospho-ERK antibody (#4370s) from Cell Signaling (Beverly MA) at 1:1000 dilution in 5% BSA solution. Loading control is measured by using α/β Tubulin antibody (#2148S) from Cell Signaling (Beverly MA) at 1:1000 dilution in 5% BSA solution.

**Table S1.**
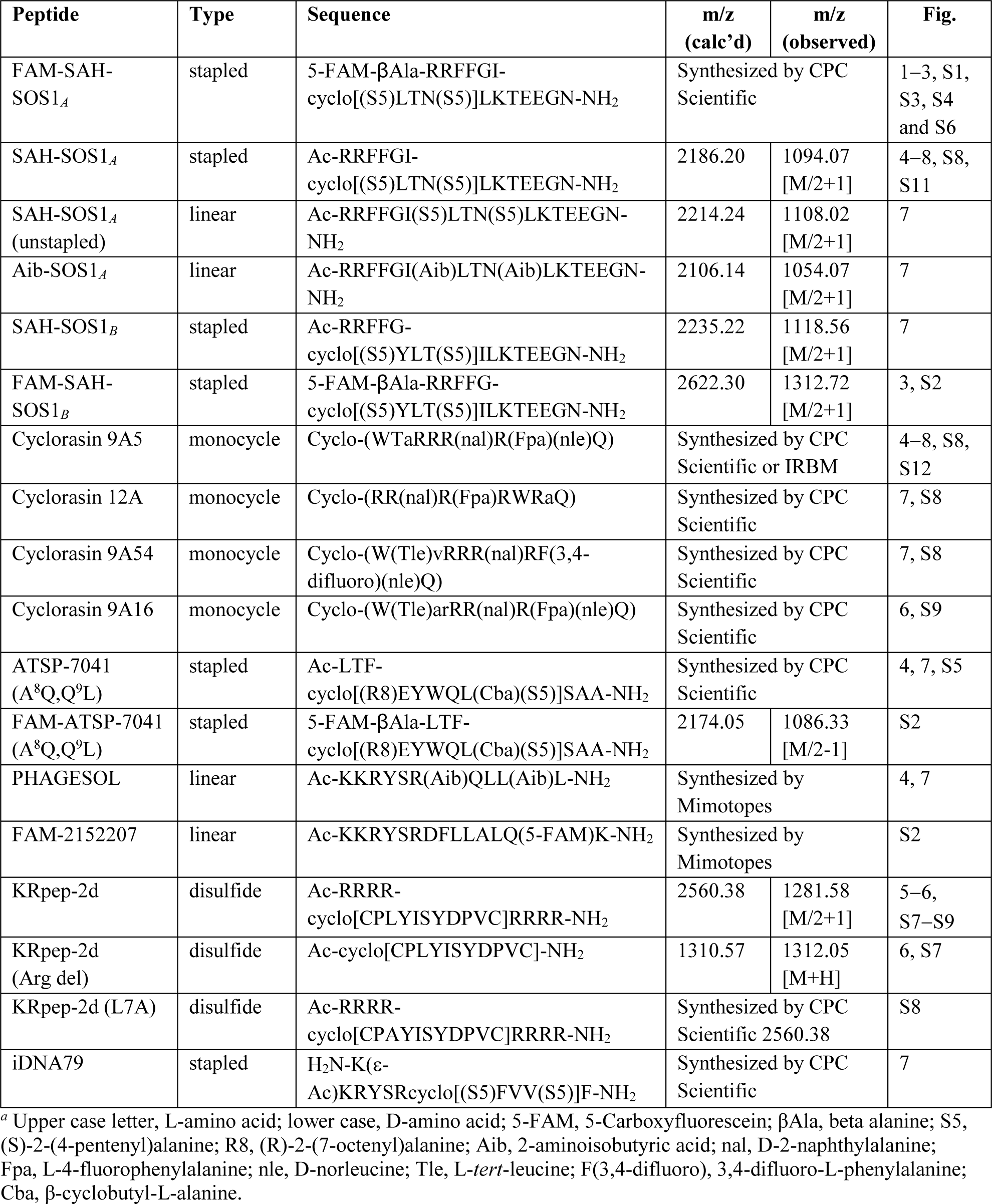
Peptides used in the studies.*^a^*

**Fig. S1.**
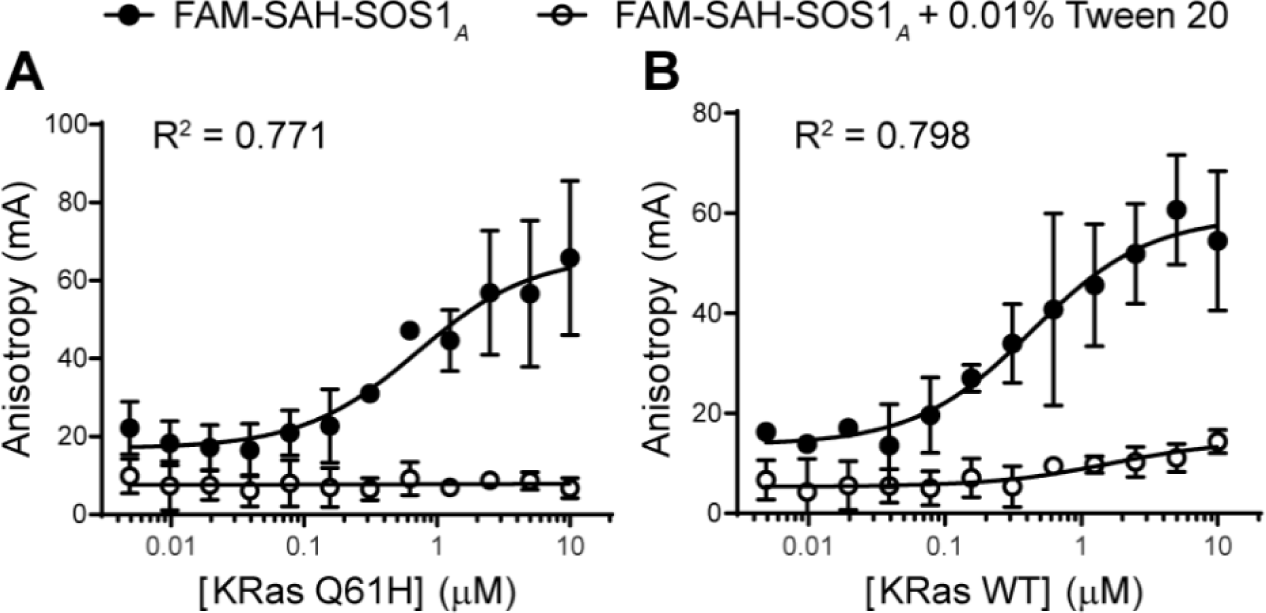
We performed the FP assay (direct binding) using FAM-SAH-SOS1*_A_* (15 nM) and measured the signals right after the mixing. We observed complete loss of binding of FAM-SAH-SOS1*_A_* to both (*A*) KRas Q61H and (*B*) KRas WT, in the presence of a small amount of detergent. Data are mean of technical triplicates ± SD. R^2^ displayed in the Fig. belongs to fitting of data from no-detergent condition.

**Fig. S2.**
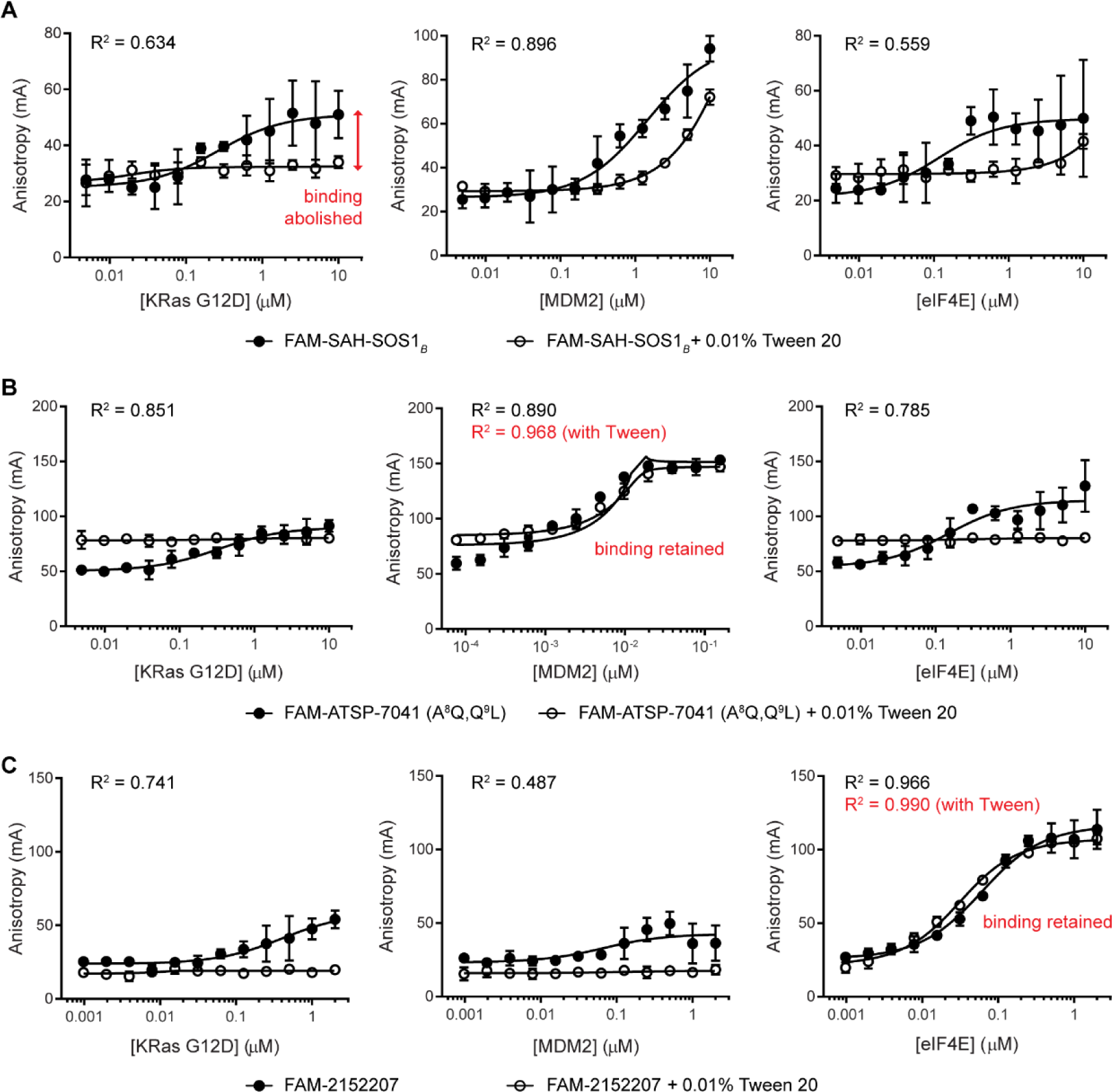
We performed the FP assay (direct binding) using FAM-labeled peptide (15 nM) and varying concentration of three distinct protein targets. FP signals are measured right after the mixing. (*A*) We observed increase in FP readouts across three different targets, KRas G12D, MDM2 and eIF4E, for FAM-SAH-SOS1*_B_* when the protein concentrations were increased. The magnitude of the increase is less than that of FAM-SAH-SOS1*_A_* (see Fig. 1). The apparent bindings were abolished upon adding of detergent in the assay buffer. (*B*) FAM-ATSP-7041 (A^8^Q, Q^9^L) could bind to MDM2 (R^2^ of fitting is 0.968) even in the presence of detergent. The result suggests that this peptide is a genuine binder for MDM2. In the absence of detergent, FAM-ATSP-7041 (A^8^Q, Q^9^L) produced non-specific FP readout regardless of protein used and the fittings are sub-optimal (R^2^ in the range of 0.785 to 0.890). (*C*) FAM-2152207 could bind to eIF4E (R^2^ of fitting is 0.990) even in the presence of detergent. The result suggests that this peptide is a genuine binder for eIF4E. In the absence of detergent, FAM-2152207 produced non-specific FP readout regardless of protein used and the fittings are sub-optimal (R^2^ in the range of 0.487 to 0.966). Data are mean of technical triplicates ± SD. R^2^ (black) displayed in the Fig. belongs to the fitting of no-detergent data (solid circle). Whereas, R^2^ (red) belongs to plus-detergent data (open circle).

**Fig. S3.**
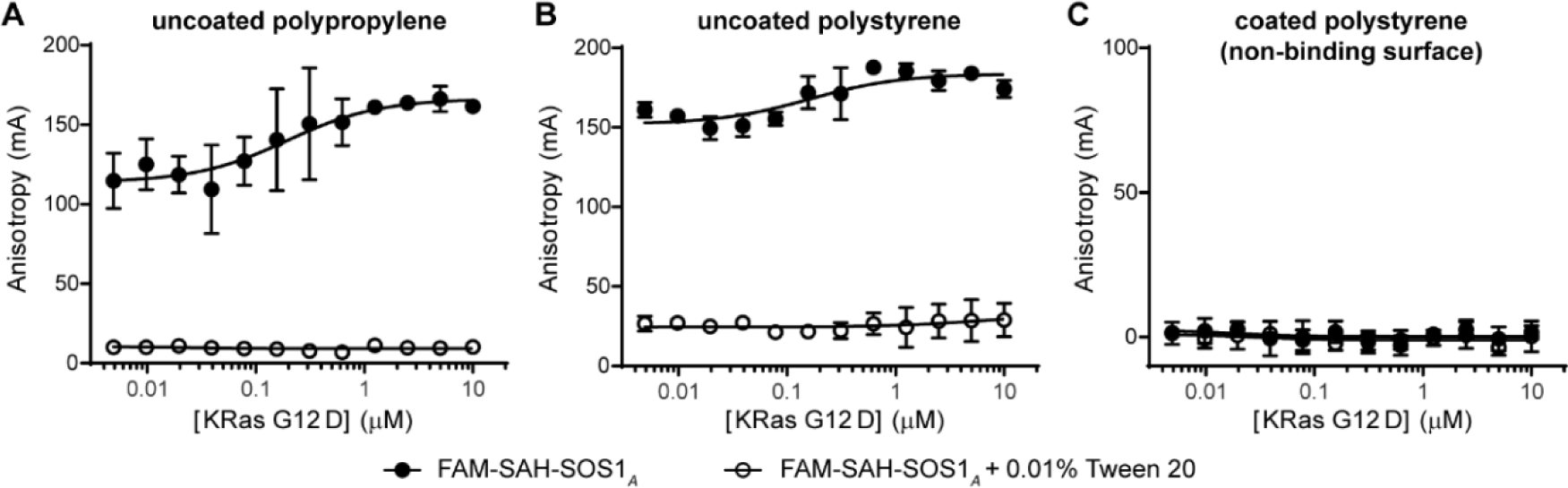
We performed the FP assay using 15 nM of FAM-SAH-SOS1*_A_* and measured the signals one hour after the mixing of the peptide and KRas G12D. Overall, the FP signals increase across all concentration tested as compared to the FP signals measured immediately after the mixing (see Fig. 2). The global signal hike was observed for assay performed on (*A*) uncoated polypropylene microplate, and (*B*) uncoated polystyrene microplate under no-detergent condition, but not on (*C*) polystyrene microplate coated with hydrophilic material (Corning NBS^TM^). The FP signal remained stable across all experiments when detergent was added (see Fig. 2). Data are mean of technical triplicates ± SD.

**Fig. S4.**
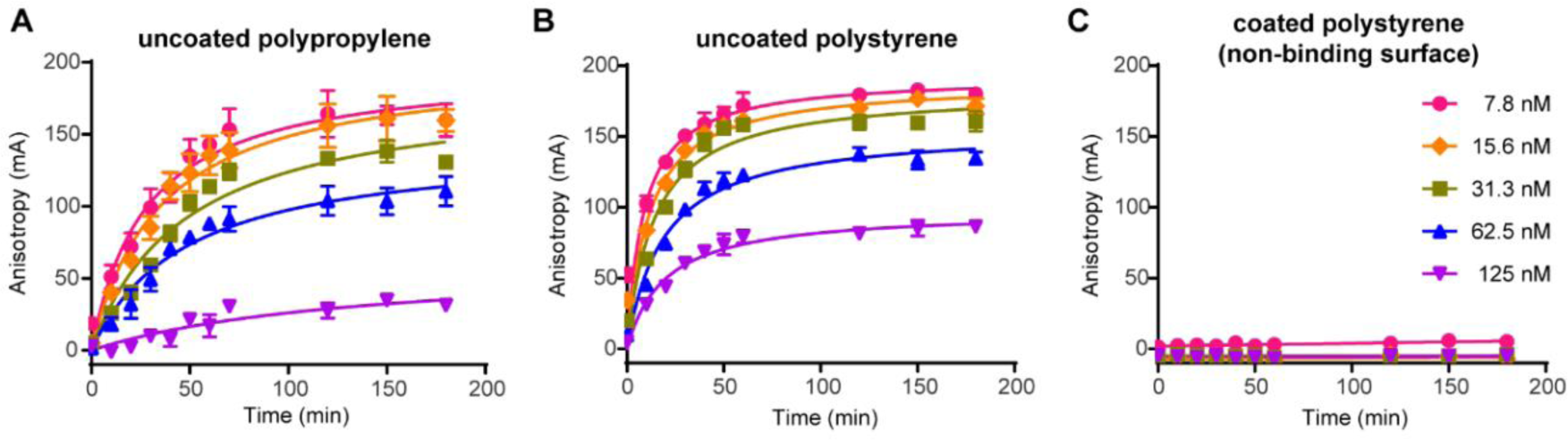
We monitored the FP signals for varying concentration of FAM-SAH-SOS1*_A_* in the absence of both KRas G12D and detergent over three hours. We observed time-dependent increase of the FP signals as soon as the peptide (diluted to appropriate concentration in polypropylene microcentrifuge tube) was added to the (*A*) uncoated polypropylene and (*B*) uncoated polystyrene microplate but not to the (*C*) polystyrene microplate coated with hydrophilic material (Corning NBS^TM^). After two hours, the FP readout on the uncoated polypropylene and polystyrene microplates approached plateau at a level which is inversely proportional to the concentration of the peptide. Intuitively, with increasing concentration, more peptides would stay in the bulk solution because of the saturation of plastic surface by the adsorbed peptide. We note that FP is based on ratiometric fluorescence output and should be less susceptible to fluorescence quenching at high probe concentration. Data are mean of technical triplicates ± SD.

**Fig. S5.**
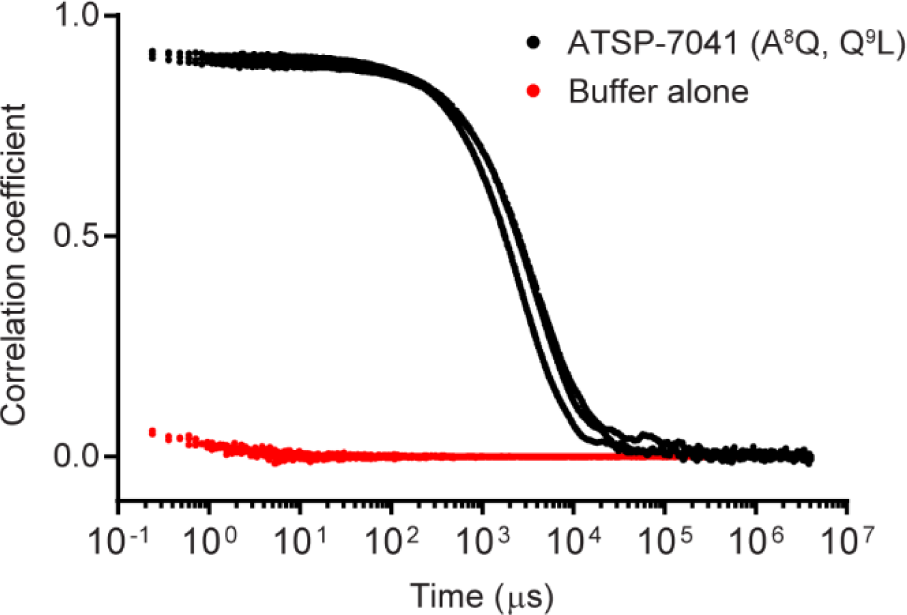
Normalized autocorrelation functions of dynamic light scattering. ATSP-7041 (A^8^Q, Q^9^L) (10 μM) underwent three independent measurements at 25 °C. The scattering intensities and the hydrodynamic diameter for ATSP-7041 (A^8^Q, Q^9^L) is 6700−8900 kcps and 560−790 nm respectively. All data are plotted.

**Fig. S6.**
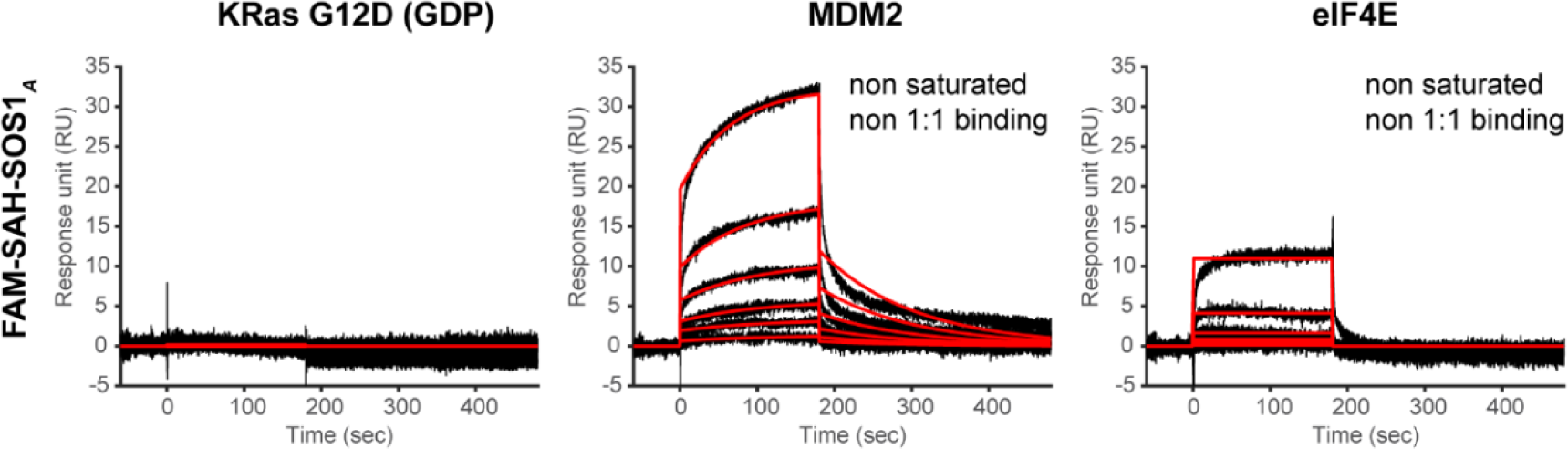
Surface plasmon resonance. We injected FAM-SAH-SOS1*_A_* over the flow cells immobilized with KRas (∼400 RU), MDM2 (∼1200 RU) and eIF4E (∼1600 RU). The injections range from 31.3 nM to 1000 nM (2-fold serial dilution). FAM-SAH-SOS1*_A_* demonstrated no binding to KRas but promiscuous non 1:1 binding to MDM2 and eIF4E. Black lines depict the double-referenced sensograms; red lines depict the global fit of the data to a 1:1 binding model.

**Fig. S7.**
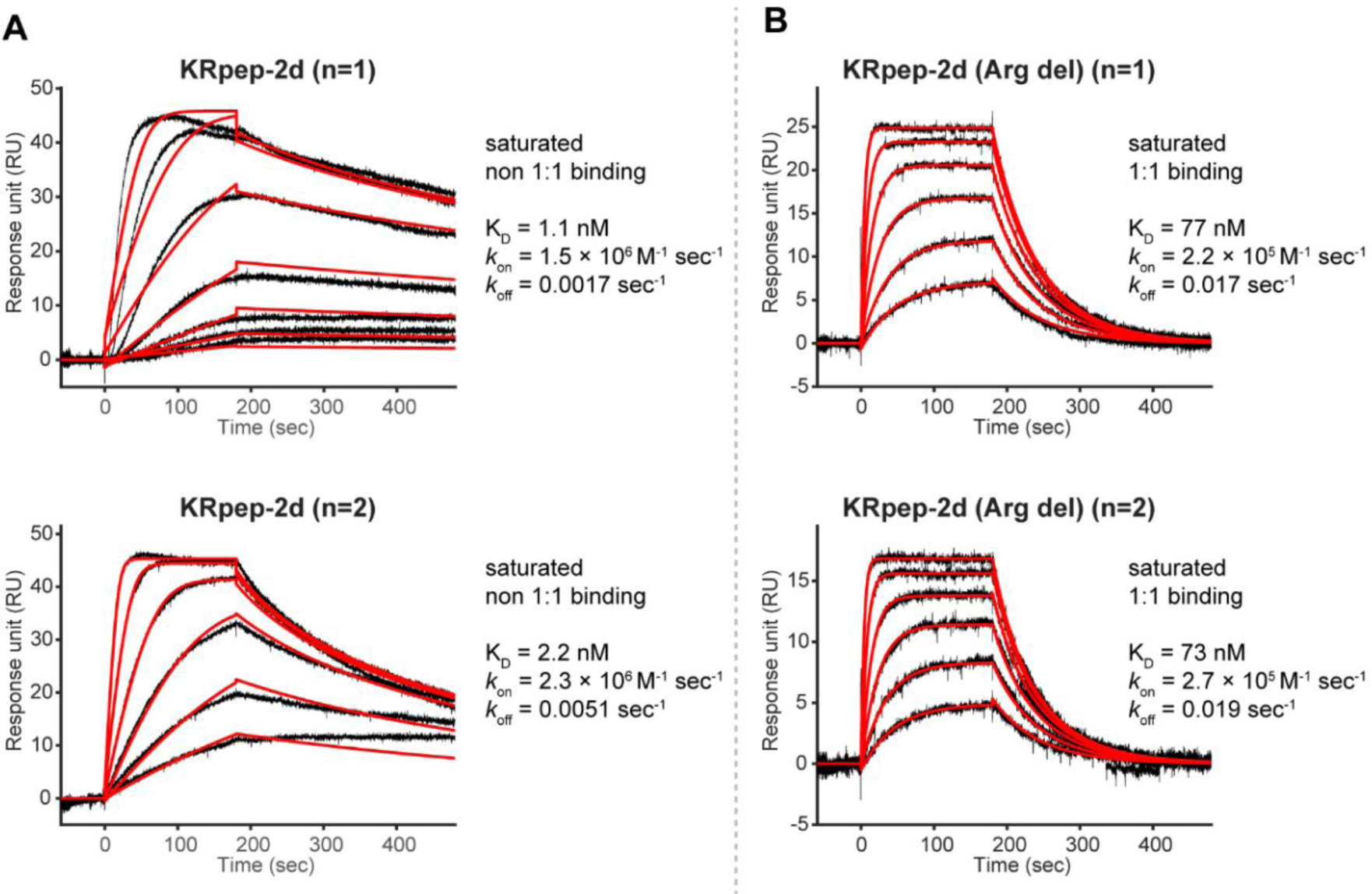
Surface plasmon resonance detected binding of positive control to GDP-loaded KRas G12D. (*A*) The top sensograms describe binding of KRpep-2d (50 nM from top, 2× dilution). The bottom sensograms describe binding of KRpep-2d (100 nM from top, 2× dilution) performed on a separate day. In both experiments, the binding approached saturation with increased peptide concentration. However, the data did not fit well to the 1:1 binding model. We note that KRpep-2d contains four arginine residues appended to both the N- and C-terminus of the sequence. We deduce that the poor fitting might derive from non-specific interaction between the peptide (+7 net charge) and the negatively charged carboxymethylated dextran coated on the gold surface of the sensor chip. (*B*) In contrast, we observed 1:1 saturated binding for KRpep-2d (Arg del), in which all the eight arginine residues flanking the sequence were removed. The injections range from 31.3 nM to 1000 nM (2-fold serial dilution). Repeating the experiment on a separate day gave similar results. The K_D,_ *k*_on_ and *k*_off_ for KRpep-2d (Arg del) is comparable with that of KRpep-2 reported in the publication (K_D_ = 51 nM, *k*_on_ = 2.3 × 10^5^ M^-1^ sec^-1^, *k*_off_ = 0.012 sec^-1^) (12). Unlike KRpep-2d, KRpep-2 only has two arginine residues flanking the sequence.

**Fig. S8.**
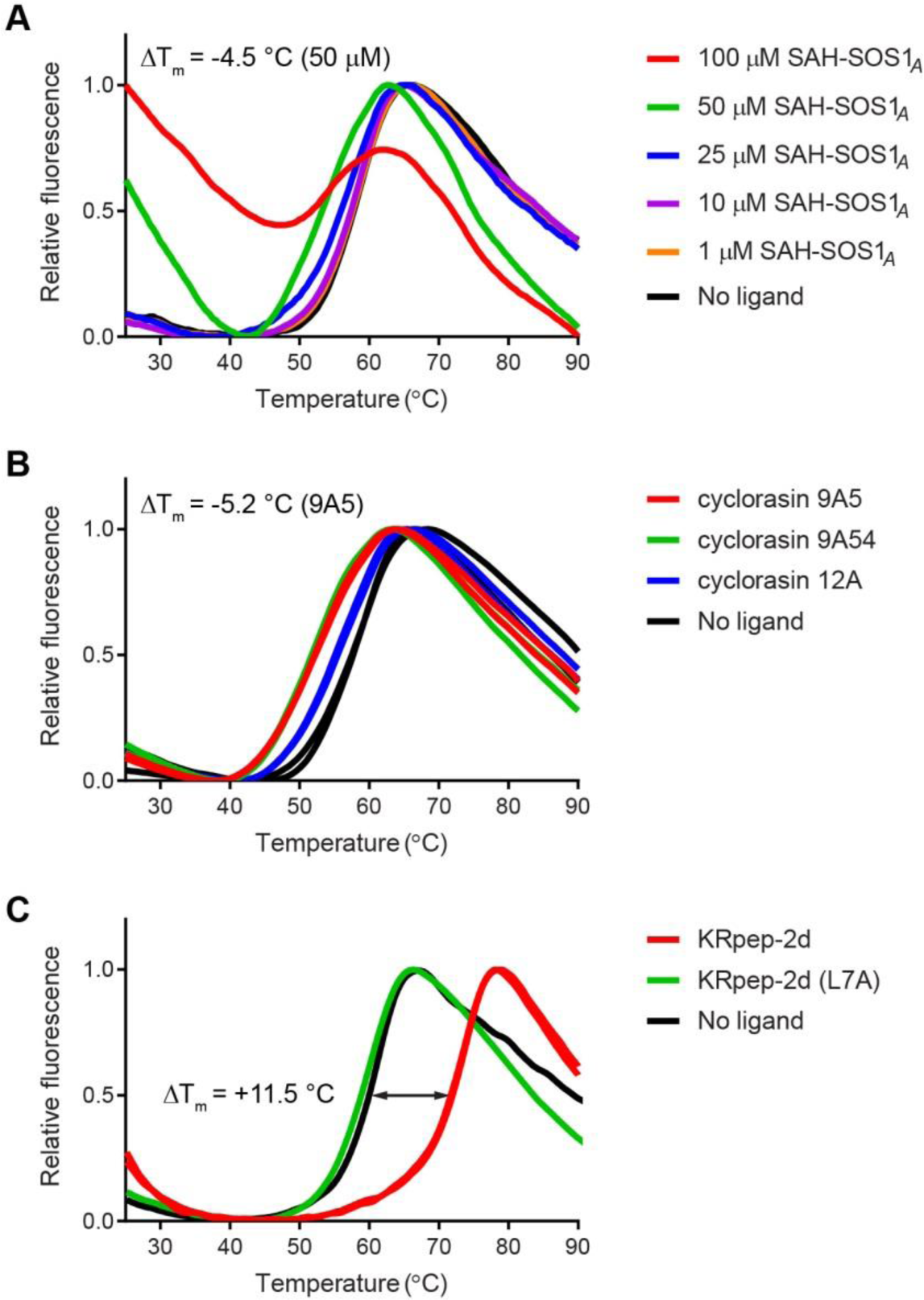
Thermal shift assay. For ease of comparison, the maximal and minimal fluorescence intensity in each data set are normalized to 1.0 and 0.0 respectively. (*A*) Melting curves of KRas G12D (10 μM) in the presence of increasing concentration of SAH-SOS1*_A_*. The shifts in protein melting temperature (ΔT_m_) are negative when compared to the no ligand control. Negative ΔT_m_ suggested destabilization of the protein by the ligand. At 50 and 100 μM of SAH-SOS1*_A_*, we observed high initial fluorescence at the starting temperature, probably due to the interaction of the fluorescent probe (SYPRO Orange) with the aggregated peptide. These signals (the non-specific interaction) decrease gradually as the temperature increases. (*B*) Melting curves of KRas G12D (10 μM) in the presence of cyclorasin 9A5 (100 μM), cyclorasin 9A54 (100 μM) or cyclorasin12A (100 μM). All three peptides induce negative ΔT_m_, although the extent is greater for cyclorasin 9A5 and cyclorasin 9A54. (*C*) Melting curves of KRas G12D (5 μM) in the presence of 500 μM KRpep-2d or 500 μM KRpep-2d (L7A). KRpep-2d showed large positive ΔT_m_ consistent with ligand-induced stabilization. Importantly, KRpep-2d (L7A) with a mutated key interacting residue (Leu to Ala) exhibited no effect on the ΔT_m_. Such results implied that the negative ΔT_m_ induced by SAH-SOS1*_A_*, cyclorasin 9A5 and cyclorasin 9A54 could be due to the destabilization of protein by the presence of colloidal aggregates formed by these peptides. In panel (*B*) and (*C*), data from technical duplicates are plotted.

**Fig. S9.**
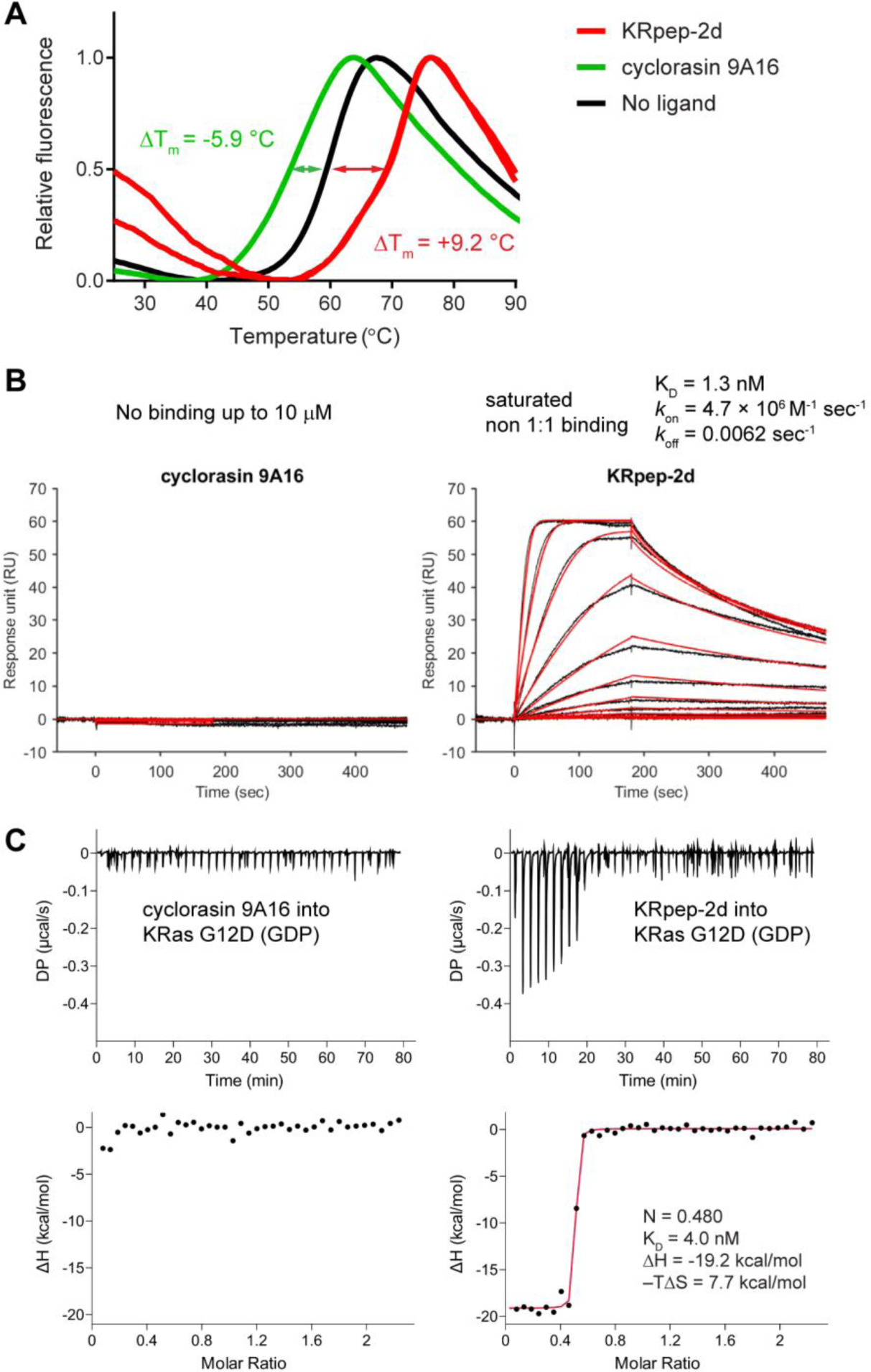
Characterization of the binding of cyclorasin 9A16 to GDP-loaded KRas G12D using various biophysical assays. (*A*) Melting curves of KRas G12D (10 μM) in the presence of either cyclorasin 9A16 (100 μM), KRpep-2d (100 μM) as positive control, or DMSO as no-ligand control. Negative ΔT_m_ from cyclorasin 9A16 suggested destabilization of the protein by the peptide. Whereas, KRpep-2d showed large positive ΔT_m_ consistent with ligand-induced stabilization. Data from technical duplicates are plotted. (*B*) Surface plasmon resonance confirmed that cyclorasin 9A16 does not bind to KRas when the peptide was tested up to 10 µM. Whereas, the positive control, KRpep-2d (50 nM from top, 2× dilution) demonstrated saturated binding to KRas. SPR experiment of KRpep-2d was performed after that of cyclorasin 9A16 using the same sensor chip. (*C*) Isothermal titration calorimetry further confirmed that cyclorasin 9A16 is a non-binder for KRas. The ITC experiment was validated subsequently on the same day using the same batch of protein and KRpep-2d which demonstrated tight binding to KRas.

**Fig S10.**
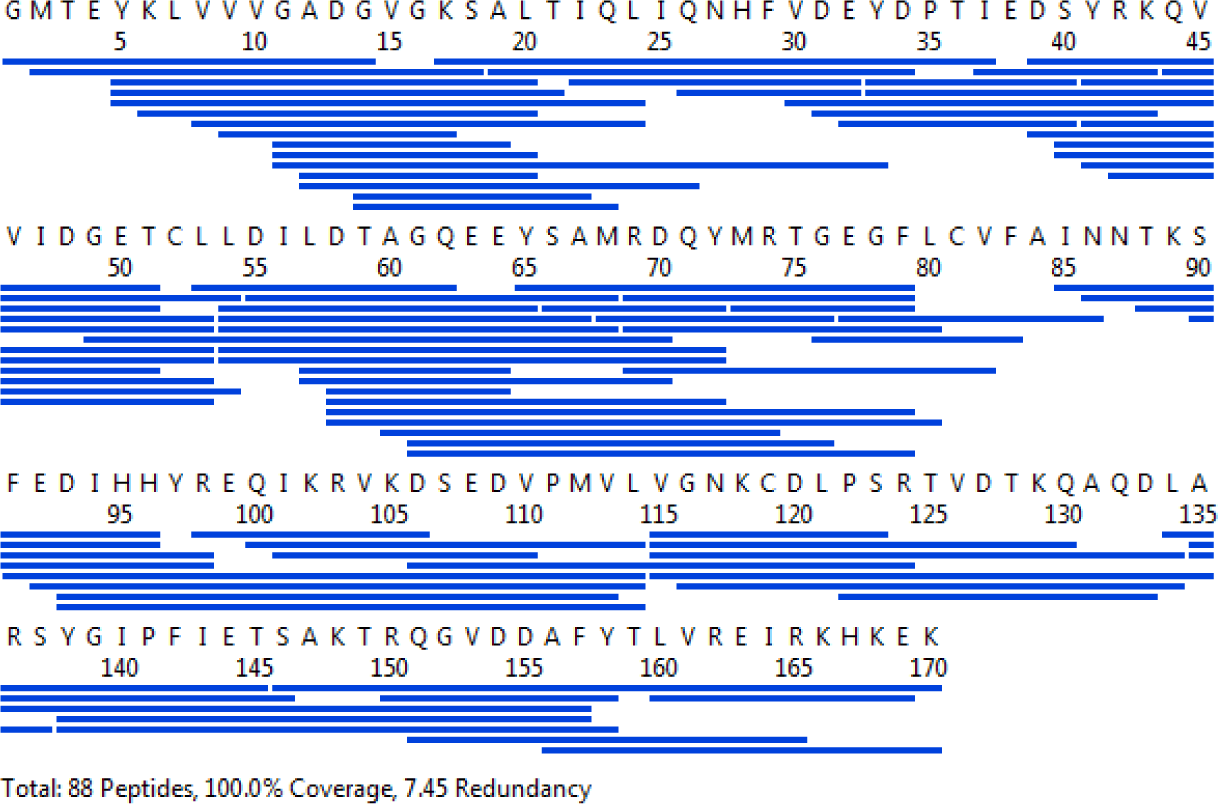
Sequence coverage map of KRas G12D from HDX-MS. Each blue solid line denotes a pepsin-proteolyzed peptide obtained. A total of 88 peptides were obtained with primary sequence coverage of 100%.

**Fig. S11.**
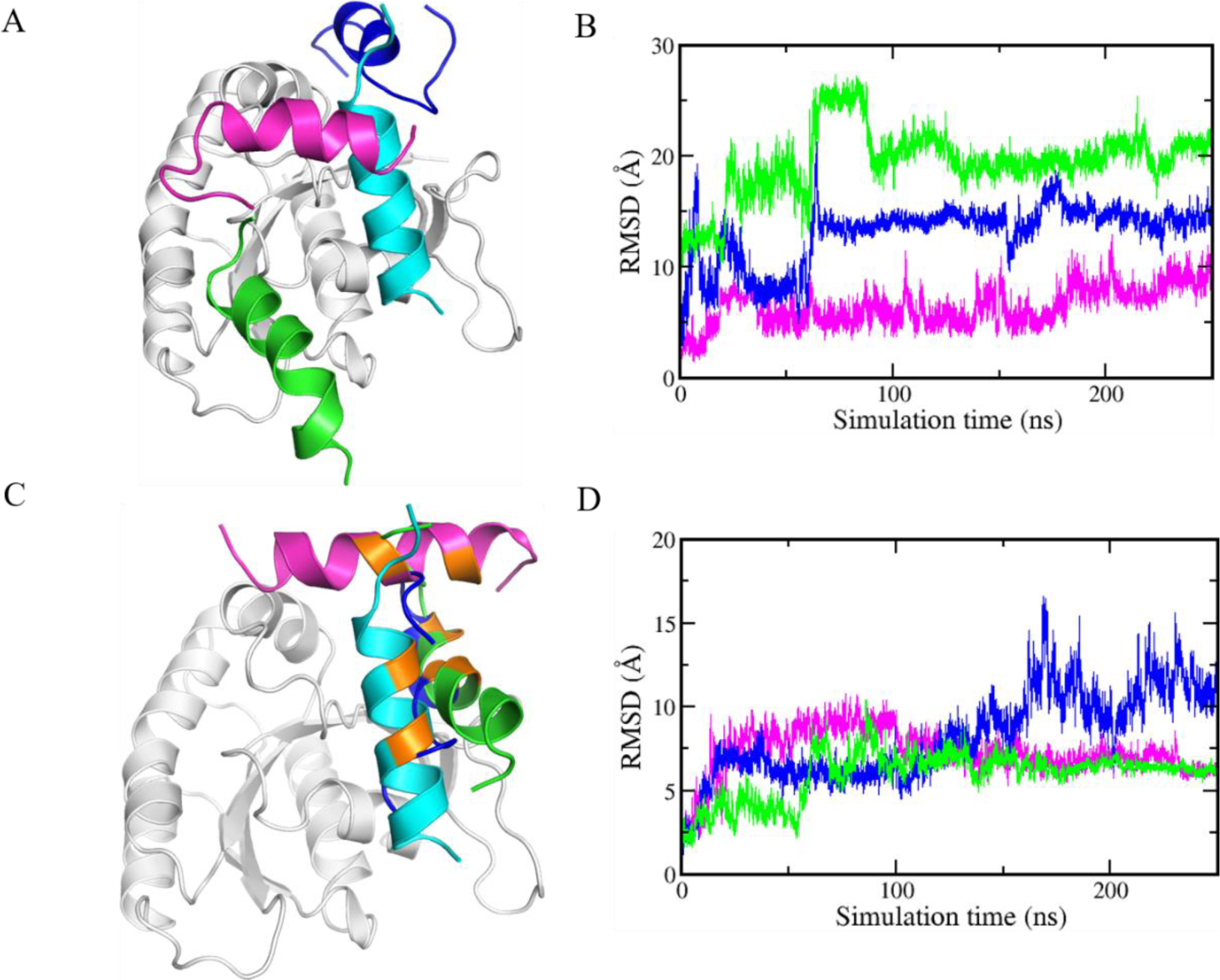
Results of MD simulations of the peptides bound to KRAS (in grey cartoon): The SOS peptides (residues 929-944 taken from the crystal structure of HRAS complexed with the SOS protein, 1NVU.pdb) are unstably bound to KRAS regardless of whether it is in apo or GDP/GTP bound states as is clear from the movement away from the initial crystallographic position shown in (*A*) Cyan coloured, starting crystallographic conformation; the peptides move away to different positions and example conformations are shown in Blue (apo KRAS); Green (GTP-bound KRAS); Magenta (GDP-bound KRAS). This can again be seen in the root mean squared deviations of the peptides during the MD simulations with respect to the starting crystallographic position (*B*). The same instability is seen for the stapled peptide SAH-SOS1*_A_*: structural deviations from the starting pose are shown in (*C*) and root mean squared deviations are shown in (*D*); colouring scheme is the same as in (*A*) and (*B*).

**Fig. S12.**
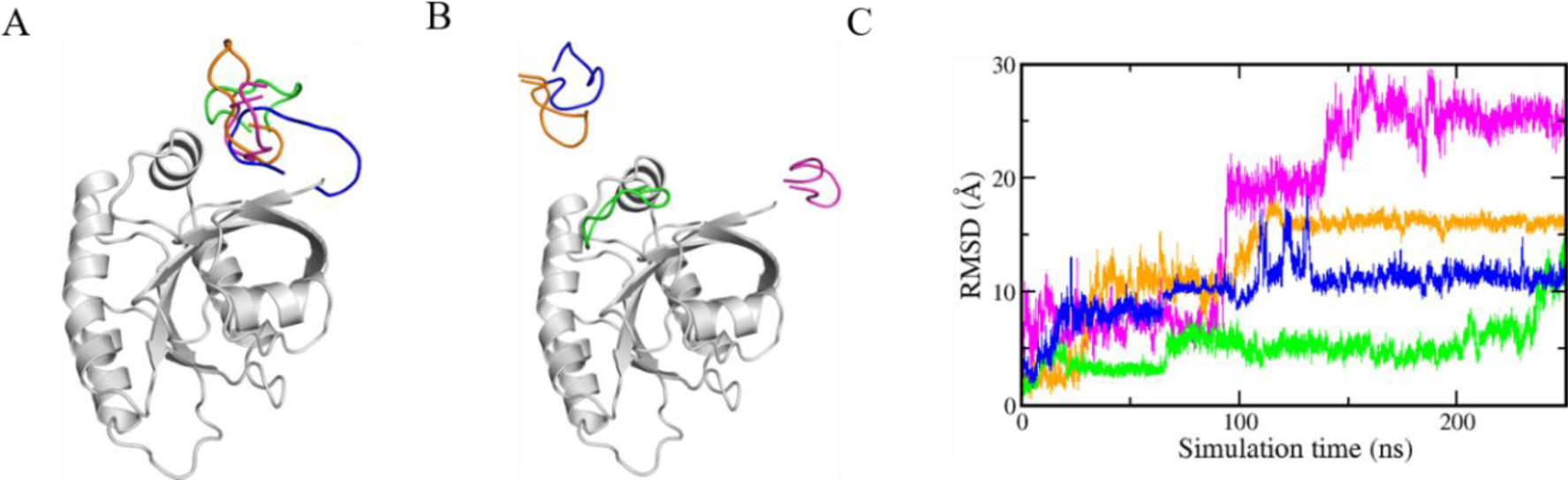
The instability of the cyclorasin 9A5 in its binding to the GTP bound form of KRAS; KRAS is shown in grey cartoon; multiple conformations of cyclorasin 9A5 were docked to KRAS; starting states shown in (*A*). All the peptides drifted away during the MD simulations (*B*). The colours of cyclorasin 9A5 correspond to the root mean squared deviations of the peptides relative to their starting structures during the MD simulations (*C*).

